# Inhibition of mutant IDH1 promotes cycling of acute myeloid leukemia stem cells

**DOI:** 10.1101/2022.04.06.487420

**Authors:** Emily Gruber, Joan So, Alexander C. Lewis, Rheana Franich, Rachel Cole, Luciano G. Martelotto, Amy J. Rogers, Eva Vidacs, Peter Fraser, Kym Stanley, Lisa Jones, Anna Trigos, Niko Thio, Jason Li, Brandon Nicolay, Scott Daigle, Adriana E. Tron, Marc L. Hyer, Jake Shortt, Ricky W. Johnstone, Lev M. Kats

## Abstract

Acute myeloid leukemias (AML) are comprised of multiple cell types with distinct capabilities to propagate the disease and resist therapy. Approximately 20% of AML patients carry gain-of-function mutations in IDH1 or IDH2 that result in over-production of the onco-metabolite 2-HG. Although IDH inhibitors can induce complete morphological remission, almost all patients eventually relapse. Analysis of clinical samples suggests that a population of IDH mutant cells is able to persist during treatment eventually acquiring 2-HG independence and drug resistance. Herein we characterized the molecular and cellular responses to the clinical IDH1 inhibitor AG-120 at high resolution using a novel multi-allelic mouse model of IDH1 mutant AML. We demonstrate that AG-120 exerts cell type-dependent effects on leukemic cells promoting delayed disease regression. Although IDH1 inhibition alone was not able to fully eradicate the disease, we uncovered that it increases cycling of rare leukemic stem cells and triggers transcriptional upregulation of the pyrimidine salvage pathway. Accordingly, AG-120 sensitized IDH1 mutant AML to azacitidine with the combination of AG-120 and azacitidine showing vastly improved efficacy *in vivo.* Our data highlight the impact of non-genetic heterogeneity on treatment response and provide mechanistic rationale for a drug combination that is being tested in clinical trials.

**STATEMENT OF SIGNIFICANCE:** Inhibition of mutant IDH1 in AML is insufficient to eliminate the disease but promotes proliferation of quiescent leukemic stem cells. Our data provide a mechanistic explanation for the observed synergy between IDH inhibitors and azacitidine and suggest that IDH inhibitors may also synergize with other drugs that preferentially target actively dividing cells.

## INTRODUCTION

Acute myeloid leukemia (AML) is a low survival cancer with a 5-year overall survival rate that still languishes below 30%. As is the case with normal hematopoiesis, AML cells are organized in a developmental hierarchy that partially recapitulates myeloid differentiation. At the apex of the hierarchy are rare leukemic stem cells (LSCs) that share many properties of hematopoietic stem cells (HSCs) including similar gene expression programs and capacity for self-renewal. LSCs in turn give rise to “bulk” tumor cells that have limited proliferative capacity but are not fully differentiated and exert pathologic effects on their microenvironment (Bonnet and Dick, 1997; van Galen et al., 2019). This non-genetic heterogeneity is a significant challenge for therapeutic targeting and the inability to fully eliminate LSCs has been linked with poor outcomes in model systems and AML patients (Fong et al., 2015; Ng et al., 2016).

Recurrent mutations in *IDH1* and *IDH2* genes were initially uncovered by cancer genome sequencing projects more than a decade ago and occur in approximately 20% of AML as well as a range of other cancers (Mardis et al., 2009; Parsons et al., 2008). Mutations are typically heterozygous and affect arginine residues within the enzymatic active site, most commonly R132 of IDH1 or R140/R172 of IDH2 (Papaemmanuil et al., 2016). Mutant IDH proteins possess a neomorphic enzymatic activity reducing α-ketoglutarate (α-KG) to the rare but structurally similar metabolite D-2-hydroxyglutarate (2-HG) (Dang et al., 2009). 2-HG accumulates to millimolar concentrations in IDH mutant cells and has been shown to deregulate α-KG-dependent pathways in a cell context-dependent manner. Critical cancerassociated proteins that are affected by 2-HG include the TET family of enzymes that mediate DNA demethylation; the JmjC domain containing histone demethylases; propyl hydroxylases that regulate hypoxic signaling and collagen maturation; and the BCAT1/2 transaminases that catalyze the synthesis of branched-chain amino acids (Figueroa et al., 2010; Koivunen et al., 2012; McBrayer et al., 2018; Sasaki et al., 2012a; Xu et al., 2011).

Multiple studies have confirmed mutant IDH1/2 as bona-fide oncogenes. IDH mutations confer anchorage and cytokine-independent growth in various cell types *in vitro* and co-operate with additional genetic lesions to initiate cancer *in vivo* (Chen et al., 2013; Kats et al., 2014; Losman et al., 2013; Lu et al., 2012). Extensive pre-clinical evidence supports the notion that mutant IDH proteins are highly promising drug targets in AML. First and foremost, IDH mutant cells demonstrate an oncogene-addicted phenotype, at least in some genetic settings. Pharmacological inhibition of 2-HG production or genetic depletion of mutant IDH can abrogate *in vitro* surrogates of transformation such as colony formation and cytokine independence, and critically reduce disease burden *in vivo* in mouse models (Kats et al., 2014; 2017; Losman et al., 2013; Shih et al., 2017). As mutant IDH enzymes possess a neomorphic biochemical activity, this putatively provides a wide therapeutic window for targeting malignant cells without affecting normal tissues. Additionally, as 2-HG is readily detectable in the serum of AML patients, it serves as a useful biomarker of pharmacodynamic efficacy (Amatangelo et al., 2017). Finally, deep sequencing and single cell studies designed to detect clonal architecture have shown that IDH mutation is an early event in leukemia evolution and is present in all or the vast majority of AML cells (Miles et al., 2020; Papaemmanuil et al., 2016; Quek et al., 2018).

In the past 5 years inhibitors of mutant IDH proteins have entered clinical use, demonstrating ∼40% overall response rate in relapsed refractory AML as single agents (Amatangelo et al., 2017; DiNardo et al., 2018). Responses to these drugs are broadly associated with differentiation of leukemic blasts, consistent with the known effect of 2-HG on myeloid development (Kats et al., 2014; Sasaki et al., 2012b). Notably however, the cellular and molecular details of the impact of IDH inhibition on the malignant hierarchy remain incompletely understood. Moreover, although some patients achieve complete morphological remission, almost all eventually relapse with the majority of cases retaining expression of the mutant IDH allele and effective blockade of 2-HG production by the inhibitor (Choe et al., 2020; Quek et al., 2018). These observations suggest that a small population of IDH mutant malignant cells with the properties of LSCs can persist during treatment and serve as the reservoir for relapse.

Herein, we developed a novel inducible mouse model of IDH1 mutant AML that enabled us to study the response of different malignant cell types to the IDH1 inhibitor AG120 (Ivosidenib). By leveraging single cell transcriptomics, we provide insights into nongenetic heterogeneity in AML and identify a novel pathway to target LSCs.

## RESULTS

### Development of a multigenic AML model with inducible expression of IDH1^R132H^

To explore the role of mutant IDH1 in leukemia maintenance *in vivo* we developed a novel mouse model termed I1DN using a methodology that we had previously applied to study mutant IDH2 (Kats et al., 2017). I1DN tumors were generated by co-transduction of fetal liverderived hematopoietic stem and progenitor cells (HSPCs) with three retroviral vectors encoding a doxycycline (dox)-regulated *IDH1^R132H^* allele linked to a dsRed fluorescent reporter, and constitutive *DNMT3A^R882H^* and *NRAS^G12D^* alleles linked to GFP and a tetracycline transactivator (tTA), respectively (Figure 1A). Importantly, this triple mutant combination is amongst the most prevalent in IDH1 mutant AML and recent single cell DNA sequencing analyses have confirmed that the mutations co-occur in individual leukemic cells (Cancer Genome Atlas Research Network et al., 2013; Miles et al., 2020; Morita et al., 2020; Papaemmanuil et al., 2016). Primary recipient mice transplanted with co-transduced HSPCs succumbed to a lethal myeloid leukemia characterized by splenomegaly and leukocytosis within 5-9 weeks (Figure 1B-D). Consistent with functional co-operation between the 3 oncogenes, donor-derived cells in the peripheral blood, spleen and bone marrow were almost exclusively GFP^+^dsRed^+^, even though this population represented only a fraction of the cells that were transplanted. Serial transplant of leukemic cells from primary recipients into syngeneic or immune-compromised NOD scid gamma (NSG) mice recapitulated disease however with shorter latency (Figures 1B and S1A). The expression of all three mutant proteins was further confirmed by western blotting and LC-MS analysis of plasma from moribund animals revealed the expected accumulation of 2-HG that was proportional to disease burden (Figures S1B-C). Administration of dox-supplemented food and water reduced expression of dsRed-linked IDH1^R132H^ in the bone marrow and spleen to near base-line levels within 5 days, whereas expression of GFP-linked DNMT3A^R882H^ and NRAS^G12D^ were maintained (Figures S1D-F).

**Figure 1:**
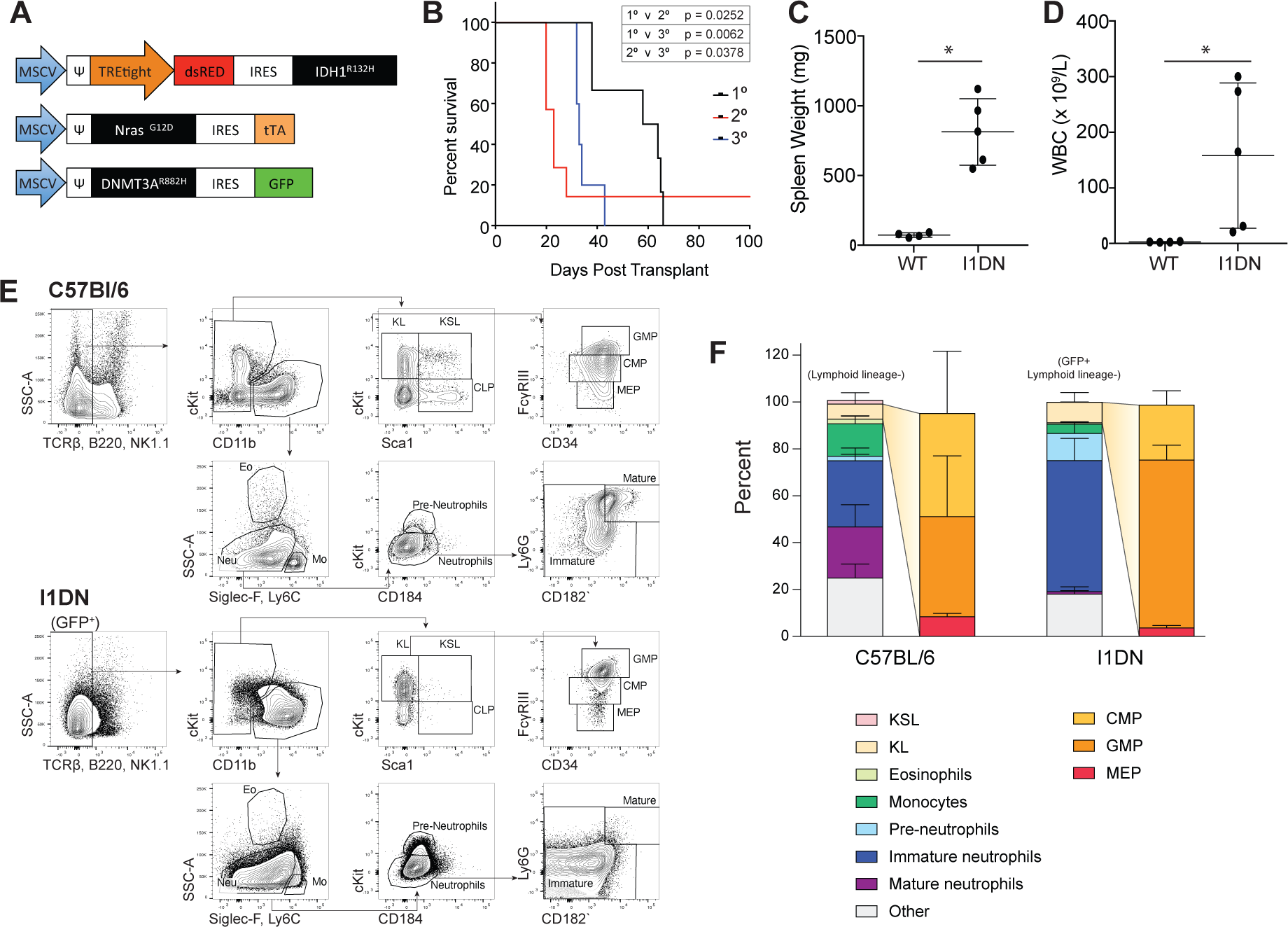
Co-expression of *IDH1^R132H^*, *DNMT3A^R882H^* and *Nras^G12D^* drives AML *in vivo*. **(A)** Schematic of retroviral constructs used to generate the I1DN model. **(B)** Kaplan-Meier survival curve of mice transplanted with transduced hematopoietic stem and progenitor cells (HSPCs, 1° recipients), or with leukemic cells from spleen or bone marrow of moribund mice (2° and 3° recipients)(n = 5-6 mice/transplant). **(C-D)** Spleen weight **(C)** and peripheral white blood cell counts **(D)** of 1° I1DN recipients or control C57BL/6 wild-type mice (n = 4-5; data represented as mean ± SD; P value was calculated by two-tailed, non-parametric student’s ttest * P < 0.05). **(E)** Representative FACS plots of myeloid progenitors and mature myeloid lineages present in the bone marrow of C57BL/6 mice and leukaemic (GFP+) cells from moribund I1DN recipients. **(F)** Bar graph of proportion of leukaemic cell types from the bone marrow of moribund IDN transplanted mice.

Dysregulated differentiation is a classical feature of AML with leukemic cells existing in a truncated developmental hierarchy dominated by an expansion of immature cells (Quek et al., 2018; van Galen et al., 2019). We used a comprehensive panel of cell surface markers and flow cytometry to quantify hematopoietic cell subsets within our model at advanced stage disease. A small proportion (5-15%) of I1DN cells in the bone marrow and spleens of diseased mice expressed the stem-progenitor marker cKit, whereas the majority expressed the myeloid lineage marker CD11b (Figure 1E-F and S1G). The cKit^+^ population primarily encompassed cells with a committed myeloid progenitor-like immunophenotype and contained a mixture of common myeloid progenitors (CMPs; Lin^-^Sca1^-^cKit^+^CD34^+^FcgR^int^) and granulocytemacrophage progenitors (GMPs; Lin^-^Sca1^-^cKit^+^CD34^+^FcgR^high^). The CD11b^+^ population was predominantly marked by immature neutrophils with a striking block of differentiation from immature (Ly6G^-^CD182^-^) to mature (Ly6G^+^CD182^+^) neutrophils apparent in both organs (Figure 1E-F and S1G) (Evrard et al., 2018). We performed a limiting dilution transplantation assay using the two most immature I1DN cell-types, CMPs and GMPs (Figures S1H-J). Mice transplanted with sorted CMPs were the first to succumb to AML, with the disease phenotype indistinguishable from mice that were transplanted with unfractionated I1DN cells. There was a strong inverse correlation between the number of cells injected and disease latency. 800 CMPs was sufficient to initiate leukemia with 100% penetrance, whereas none of the mice injected with 800 GMPs developed disease, indicating at least a 10-fold enrichment in LSC frequency within the CMP compartment. In summary, we have developed a novel model of mutant IDH1 AML characterized by aberrant progenitor self-renewal and a block of myeloid differentiation at the immature neutrophil stage.

### Inhibition of mutant IDH1 promotes AML differentiation and prolongs survival

Small molecule inhibitors of mutant IDH proteins have demonstrated efficacy in model systems and clinical trials (DiNardo et al., 2018; Kats et al., 2017; Shih et al., 2017; Stein et al., 2017), but the molecular and cellular events downstream of 2-HG depletion remain incompletely understood. We used the I1DN model to assess the impact of IDH1 targeting *in vivo* and compared pharmacological inhibition by the clinical IDH1 inhibitor AG-120 (a.k.a. Ivosidenib) to genetic depletion of the IDH1^R132H^ transgene. We first confirmed that AG-120 therapy reduced 2-HG in the plasma and bone marrow of leukemic mice, as expected (Figure S2A). We then treated a cohort of I1DN-engrafted recipients with vehicle, AG-120 or dox and monitored disease progression and survival (Figures 2A and S2B-C). In the vehicle-treated group the rapid increase in peripheral blood tumor burden was accompanied by the development of severe thrombocytopenia, with all animals requiring euthanasia due to AML by day 38 post-transplant. By comparison, disease progression in AG-120 and dox-treated animals was arrested following 2-3 weeks of treatment and although I1DN cells continued to be detectable for the duration of the study, platelet counts normalized. 10/11 mice in the AG120 group and 12/12 mice in the dox group, survived until day 42, at which point treatment was withdrawn to determine whether the disease relapsed. Surprisingly, tumor burden remained stable although most animals developed a graft vs host disease-like condition (likely caused by co-transfer of a small number of T-cells, S2D and Table S1) and had to be withdrawn from the study, precluding long term analysis. We did however, identify at least two bona-fide relapses in the AG-120 group (#13 and #23) characterized by rapid expansion of I1DN cells and coincidental onset of thrombocytopenia. Taken together, our findings demonstrate that IDH1 inhibition is effective *in vivo* against IDH1 mutant AML and confers a significant survival benefit in treated animals but does not fully eliminate the disease.

**Figure 2:**
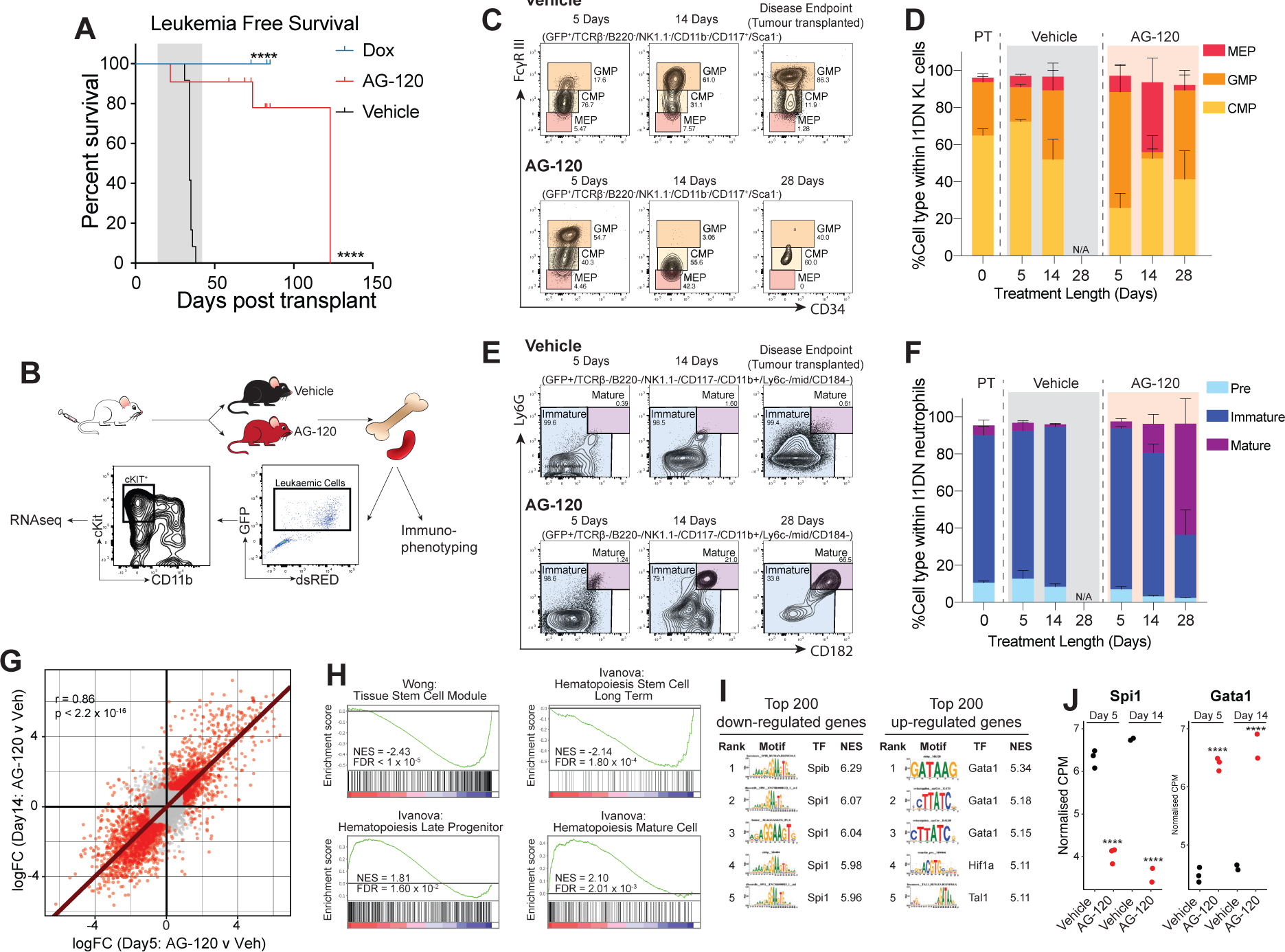
AG-120 promotes differentiation and delayed disease regression *in vivo* by depleting 2-HG. **(A)** Kaplan-Meier progression-free survival curve of I1DN tumor bearing mice treated with AG-120, dox or vehicle. Grey shading denotes treatment. Mice that succumbed to graft vs host disease were censored (n = 11-12 mice/group; P value was calculated by log-rank test ****P < 0.0001). **(B)** Schematic of experimental design. **(C-E)** Immunophenotype of I1DN leukemic cells in the bone marrow of mice treated with AG-120 or vehicle for the indicated time period. Representative FACS plots **(C and E)** and quantification **(D and F)** of the progenitor **(C-D)** and mature **(E-F)** leukemic compartments is shown. (n = 3-4 mice/group/time point; data represented as mean ± SD). **(G-J)** RNAseq analysis of sorted I1DN progenitor cells. **(G)** Correlation plot of differentially expressed genes (DEGs) induced by AG-120 following 5 and 14-days of treatment. Red circles represent significant DEGs (FDR < 0.01 and |logFC| > 1) in either comparison. Line of best fit calculated for significant DEGs in either comparison. **(H)** GSEA barcode plots showing down-regulation of HSC-associated genes and up-regulation of myeloid differentiation genes following 5 days of AG-120 treatment. **(I)** Enrichment of transcription factor binding sites identified within the proximal promoters of the top 200 AG120 up or down-regulated genes identified by RcisTarget. Ranking was determined by normalized enrichment score (NES). **(J)** Voom normalized counts of *Spi1* and *Gata1* (**** FDR < 0.0001 calculated with the limma package in R).

**Table 1:** Antibodies used for flow cytometry analysis and cell sorting.

| **Antibody** | **Source** | **Catalogue number** |
| --- | --- | --- |
| Anti-Mouse IgG/HRP | Dako | Cat# P0260 |
| Anti-Rabbit IgG/HRP | Dako | Cat# P0217 |
| B220 APC-E780 | Invitrogen | Cat# 47-0452-82 |
| β-actin | Sigma | Cat# A2228 |
| CD11b APC-E780 | Thermo Fisher | Cat# 47-0112-82 |
| CD11b Biotin | MACS Miltenyi Biotech | Cat# 130-113-233 |
| CD11b BV450 | BD Horizon | Cat# 560455 |
| CD11b BV711 | Biolegend | Cat# 560455 |
| CD16/CD32 (FCγR-III) BV786 | BD Optibuild | Cat# 740851 |
| CD16/CD32 (FCγR-III) BUV737 | BD Horizon | Cat# 565272 |
| CD34 BV421 | BD Biosciences | Cat# 562608 |
| CD115 Superbright436 | Invitrogen | Cat# 62-1152-82 |
| CD117 APC | BD Pharmingen | Cat# 553356 |
| CD117 BV650 | BD Horizon | Cat# 563553 |
| CD182 APC | Biolegend | Cat# 149312 |
| CD184 BUV395 | BD Optibuild | Cat# 740265 |
| DNMT3A | Cell Signaling | Cat# 2160s |
| IDH1 | Cell Signaling | Cat# 8137s |
| Ly-6A/E (Sca1) Pe/Cy7 | BD Pharmingen | Cat# 558162 |
| Ly-6C BV421 | Biolegend | Cat# 128032 |
| Ly-6C BV785 | Biolegend | Cat# 128041 |
| Ly-6G BV605 | Biolgened | Cat# 127639 |
| NK1.1 APC/Cy7 | BD Pharmingen | Cat# 560618 |
| NRAS<sup>G12D</sup> | Cell Signaling | Cat# 14429s |
| Siglec-F BV421 | BD Horizon | Cat# 562681 |
| Siglec-F BV786 | BD Optibuild | Cat# 740956 |
| Streptavidin APC/Cy7 | Biolegend | Cat# 405208 |
| TCRβ APC-E780 | eBioscience | Cat# 47-5961-82 |
| TCRβ E450 | eBioscience | Cat# 48-5961-82 |

To characterize the impact of IDH1 inhibition on the AML hierarchy in greater detail, we analyzed additional cohorts of tumor bearing mice that had been treated with vehicle or AG-120 for 5 days, 2 or 4 weeks (the maximum allowable time under our approved animal ethics protocol) (Figure 2B). For these and subsequent experiments we used I1DN tumors that had been depleted of lymphoid cells to reduce the incidence of graft-vs-host-disease (see Methods). AG-120 initially increased the total tumor burden in the bone marrow and spleen, and at 5 days there was a dramatic increase in spleen size and macroscopically visible regions of dense blast cell infiltrate into the medulla of the spleen of treated animals compared with controls (Figures S2E-F). The initial burst of proliferation was later followed by disease regression and notably the kinetics differed between different organs, with reduction of I1DN cells in the bone marrow and spleen preceding that in the peripheral blood. Flow cytometric analysis revealed progressive myeloid differentiation that was evident as early as 5 days after treatment initiation but continued to progress over 4 weeks (Figures 2C-F). Initially, we observed changes predominantly in the immature cKit^+^Lineage^-^ (KL) compartment (hereafter referred to as the progenitor compartment), with an increase in the GMP:CMP ratio. At later time points these immature cells were almost completely eliminated, although a small population of CMPs remained at the end of treatment. Mature (cKit^-^CD11b^+^) cells continued to accumulate, eventually acquiring the phenotype of fully differentiated neutrophils. Taken together, our data suggest that AG-120 acts at multiple levels of the AML hierarchy, altering the phenotypes of both LSCs and bulk tumor cells, leading to exhaustion and delayed tumor regression (Figure S2G).

We next analyzed the impact of IDH1 inhibition on gene expression by RNA sequencing (RNAseq) of sorted leukemic progenitor cells from I1DN tumor-bearing mice that had been treated with vehicle or AG-120 for 5 or 14 days (Figure 2B). We identified 2,207 and 2,211 differentially expressed genes (DEGs; FDR < 0.05, and |log2FC| > 0.5) following 5 and 14 days of AG-120 treatment, respectively (Table S2). Transcriptional alterations at the two time points were highly positively correlated, demonstrating that AG-120 induced changes can occur within days of treatment initiation and remain stable over time (Figure 2G and S2H).

Among the DEGs shared between 5 and 14 days of drug treatment were numerous factors with well-known roles in normal and malignant hematopoiesis. These included *Meis1*, *Sox4* and *S1pr1* which were downregulated (Supplementary Table 2). *Meis1* and *Sox4* encode stem cellassociated transcription factors that are frequently dysregulated in AML and contribute to increased self-renewal capabilities that drive uncontrolled proliferation (Wang et al., 2005; (H. Zhang et al., 2013); whereas *S1pr1* regulates egress of lineage committed HSPCs from the bone marrow into the blood (Juarez et al., 2012). On the other hand, *Prg2* and *CD177,* which have functions in mature myeloid cells, were upregulated. Unbiased analysis using gene set enrichment (GSEA) (Subramanian et al., 2005) further confirmed a strong negative enrichment of HSPC-associated gene sets and conversely, a strong positive enrichment of gene sets that govern myeloid differentiation (Figure 2H). Furthermore, using the RcisTarget algorithm we identified the Gata1/Spi1 axis as a potent driver of the AG-120 response. Gata1 and Spi1 are cross-antagonistic transcription factors that inhibit each-other’s activity while promoting their own (Hoppe et al., 2016). Gata1 and Spi1 binding motifs were highly over-represented in the most significant AG-120 up and down-regulated genes, respectively (Figure 2I). Concordantly, *Gata1* expression was increased while *Spi1* expression was reduced (Figure 2J). Thus, IDH1 inhibition alters the expression of molecular programs that control self-renewal and differentiation.

The oncometabolite 2-HG has been postulated as the primary effector through which mutant IDH proteins exert their oncogenic function (Kats et al., 2017; Koivunen et al., 2012; Losman et al., 2013). Notably, recent studies have implicated the reacquisition of 2-HG production via alternative pathways as a mechanism driving resistance to IDH inhibitor therapy in a subset of AML patients (Harding et al., 2018; Intlekofer et al., 2018; Morita et al., 2020). We compared the AG-120 transcriptional signature to a signature of mutant IDH2 inhibition that we had previously derived in an isogenic AML model driven by IDH2^R140Q^, DNMT3A^R882H^ and Nras^G12D^ (Kats et al., 2017). Using rotating gene set testing (ROAST) (Wu et al., 2010) we identified significant enrichment of DEGs that were perturbed by IDH2^R140Q^ inhibition in the AG-120 transcriptional signature (Figure S2I). We directly compared the effects of AG-120 and deinduction of the IDH1^R132H^ transgene by dox and found that both treatments elicited phenotypic differentiation of I1DN progenitors and highly similar gene expression changes by RNAseq (Figure S2J-K). Taken together, these data demonstrate that the effects of AG-120 are mediated by on target inhibition of 2-HG production.

### Resistance to IDH inhibition is associated with transcriptional reprogramming

Our initial survival study prompted us to explore whether I1DN leukemic mice that had relapsed after treatment withdrawal developed AG-120 resistant disease. The cross-sectional study shown in Figure 2B-J also included a cohort of tumor-bearing mice treated with AG-120 for 4 weeks and subsequently monitored for disease relapse. We observed delayed disease regression in the peripheral blood of these animals (Figure S3A), but none attained complete remission and I1DN cells continued to be detectable in the peripheral blood by FACS. Notably, the greatest reduction in peripheral blood tumor burden occurred after treatment withdrawal, demonstrating the lag between IDH inhibition and disease regression that likely occurs due to the time required for LSCs to differentiate. In all 4 mice disease remained stable for at least 6 weeks before relapse. The kinetics of relapse strongly suggest that I1DN cells had remained latent for several weeks after treatment cessation before reacquiring proliferative capabilities.

To gain insights into gene expression programs that may underpin the relapse in these I1DN tumors, hereafter referred to as treatment exposed (TE) leukemias, we compared the transcriptome of the I1DN TE progenitors and I1DN vehicle-treated progenitors. In multidimensional scaling (MDS) analysis I1DN TE samples formed 2 distinct clusters and were separated from vehicle-treated samples, indicating the establishment of novel transcriptional states in relapsed leukemias (Figure 3A). We combined the two samples in each cluster (TE1 and TE2) and then identified DEGs between TE clusters and vehicle-treated samples. Overall, there were 687 and 897 DEGs (FDR < 0.05 and |log2FC| > 0.5) between TE1 or TE2 and vehicle-treated, respectively, with few overlapping DEGs between TE1 and TE2 (Figure 3B, S3B and Supplemental Table 2). GSEA and gene ontology analyses revealed that TE1 was characterized by the upregulation of myeloid differentiation and inflammatory gene sets, whereas TE2 tumors upregulated stem cell and histone methylation pathways, whilst downregulating mitochondrial respiratory and oxidative phosphorylation signatures (Figure 3C-D). RcisTarget identified Spi1 and Foxo1 as the top transcription factor motifs upstream of up-regulated TE1 and TE2 DEGs, respectively (Figure S3C).

**Figure 3:**
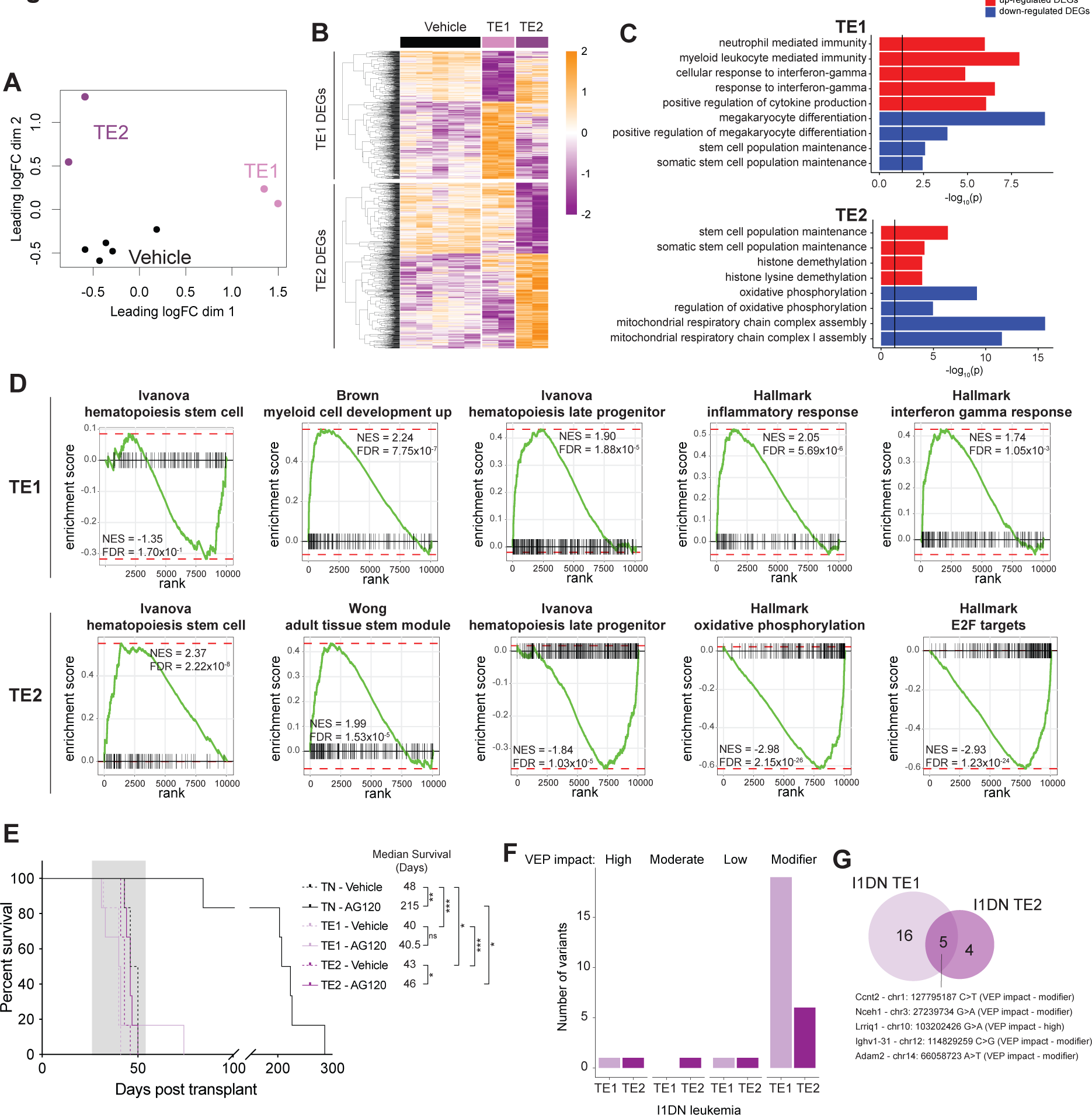
I1DN tumors resistant to AG-120 are characterized by altered gene expression. **(A-D)** RNAseq of cKit+ I1DN progenitor cells from vehicle-treated mice or AG-120 relapsed mice. **(A)** MDS plot of the top 500 most variably expressed genes. **(B)** Heatmap of TE1 and TE2 DEGs (FDR < 0.05 & |logFC| > 0.5). **(C)** GO terms of TE1 and TE2 DEGs. **(D)** GSEA pathways of TE1 and TE2 DEGs. **(E)** Kaplan-Meier survival curve of mice transplanted with I1DN TN or TE leukemias and treated with AG-120 or vehicle. (n = 6 mice/group; P value was calculated by log-rank test * P < 0.05, **P < 0.01, *** P < 0.001).TN; treatment naïve, TE; treatment exposed.

To determine whether relapsed I1DN TE tumors had acquired resistance to IDH1 inhibition, we transplanted cells from two mice (TE1; mouse#24 and TE2; mouse#23) into cohorts of secondary recipients. We also transplanted mice with I1DN treatment naïve (TN) cells. Concordant with our previous studies, AG-120 prolonged survival in the TN cohort, whereas mice transplanted with TE cells did not respond to AG-120 (Figure 3E). Of note, although the median survival of all vehicle treated cohorts differed by less than 7 days, it was slightly lower in both TE cohorts, suggesting that these tumors proliferate more rapidly at baseline.

We performed exome sequencing on a TE1 tumor, a TE2 tumor and a TN tumor (Figure 3F-G). Comparison of the mutational profiles identified only a single mutation in *Lrriq1* that was predicted to have a detrimental effect on protein function as determined by the Variant Effect Predictor functional consequence rank. *Lrriq1* expression was undetectable in I1DN progenitor cells by RNAseq. No mutations were identified in hematopoietic transcription factor or kinase signaling genes. Mutant isoform switching and gate-keeper mutations in IDH1 and 2 have been observed in a proportion of AML patients with acquired resistance to IDH inhibitors (Harding et al., 2018; Intlekofer et al., 2018). Close examination of the IDH1 and -2 loci revealed no mutations within the coding domain sequences of these genes, including the recurrently mutated hotspot codons. These findings suggest that the drug-resistant and transcriptional phenotypes of I1DN TE tumors are the result of epigenetic rather than genetic changes.

### AG-120 promotes cycling of quiescent LSCs and upregulates the pyrimidine salvage pathway

Over the past decade, it has become apparent that non-genetic drivers underpin a large proportion of functional variation in cancer, including the capacity of rare cell populations to withstand therapeutic pressure ultimately leading to drug resistance (Marine et al., 2020). To investigate the extent of heterogeneity of AG-120 transcriptional responses, we utilized the 10X Genomics 3’ single-cell RNA sequencing (scRNAseq) platform to profile gene expression from individual I1DN progenitor cells isolated from mice following 5 days of inhibitor or vehicle treatment. After filtering, we obtained high quality data for 7,708 cells with a median of 4,126 genes quantified per cell. Uniform manifold approximation and projection (UMAP) clustering identified 9 distinct transcriptional clusters (Figures S4A-B), all of which contained cells from both vehicle and AG-120 treated animals suggesting that the grouping reflected the presence of different cellular states and was not based on the molecular response to IDH inhibition (Butler et al., 2018). Correlation analysis enabled us to merge clusters that had fewer than 100 cells to highly similar larger clusters, generating 7 main progenitor groups (Figure 4A).

**Figure 4:**
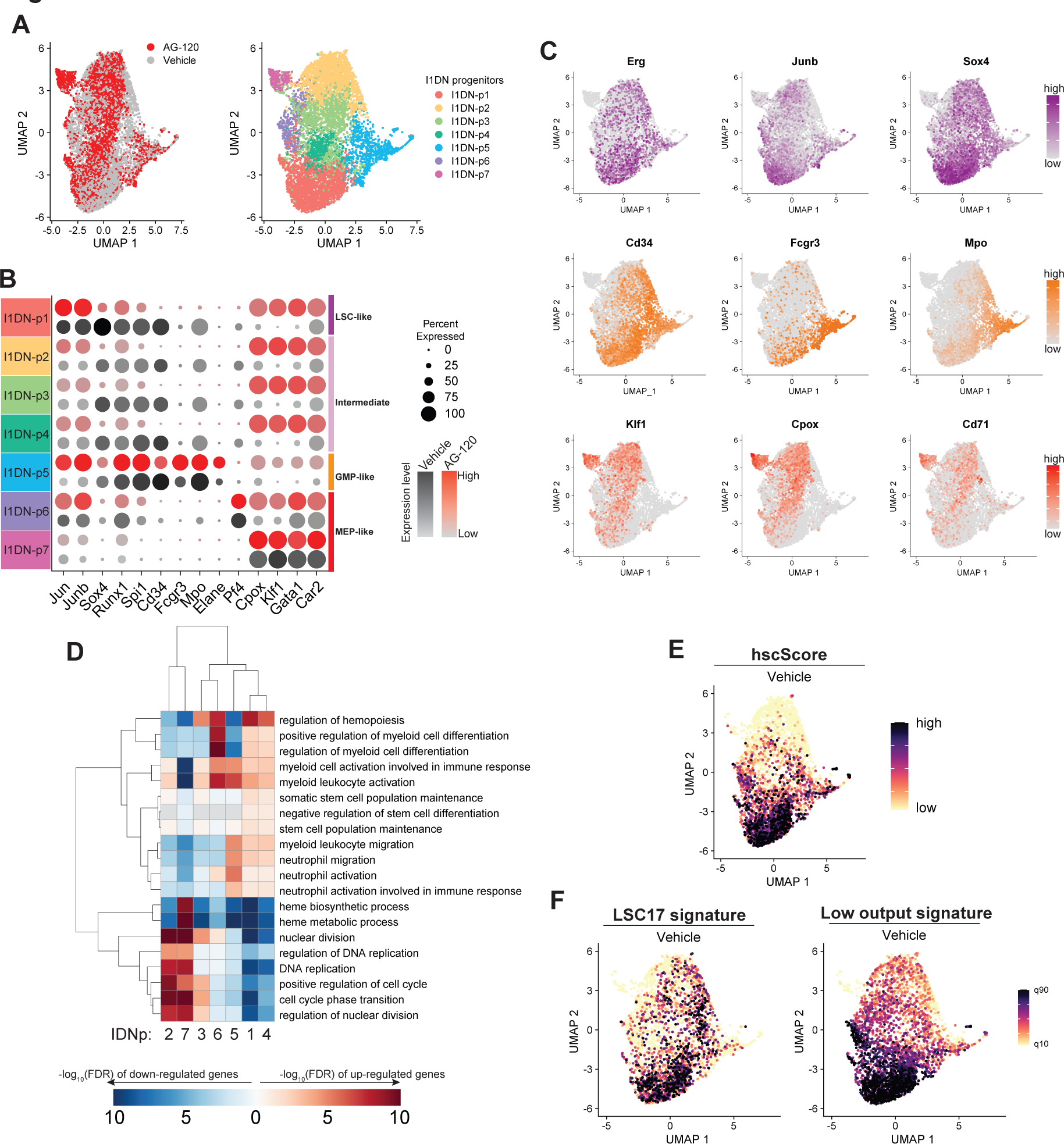
scRNAseq identifies LSCs within the I1DN progenitor population. **(A)** I1DN progenitor cells from mice treated with AG-120 or vehicle for 5 days were analyzed by scRNAseq. UMAP plot of integrated datasets labelled by treatment (left) or I1DNprogenitor (I1DN-p) group (right). **(B)** Dotplot of gene expression across IDN1p groups and treatment. **(C)** UMAP plots of highlighted genes. **(D)** Heatmap of gene ontology (GO) terms associated with genes significantly differentially expressed between the different ID1Np groups. Only cells from vehicle treated mice were used in this analysis. **(E-F)** UMAP plot of **(E)** hscScore(Hamey and Göttgens, 2019), **(F)** “low-output” HSC signature(RodriguezFraticelli et al., 2020) and LSC17 score(Ng et al., 2016). GMP – granulocyte-macrophage progenitor; MEP – megakaryocyte-erythroid progenitor; LSC – leukemia stem cell.

Each group expressed distinct patterns of well-established hematopoietic differentiation markers (Lara-Astiaso et al., 2014) that led us to annotate group 5 as GMP-like (high expression of *Fcgr3* and *Mpo*), groups 6 and 7 as MEP-like (high expression of *CD71* and *Klf1*), groups 1 as LSCs (high expression of *Erg*, *Junb* and *Sox4*) and groups 2-4 as intermediate (Figures 4B and C). Concordant with our manual annotation, cells within groups 6 and 7 upregulated expression of genes required for heme biosynthesis, whereas group 5 was marked by signatures of myeloid cell activation and groups 1-4 by various stemness associated signatures (Figure 4D). Notably, group 1 cells most closely resembled long-term HSCs as evident by their high hscScore (Hamey and Göttgens, 2019) and prominent expression of the “low-output” signature characteristic of quiescent, self-renewing, long-term repopulating HSCs recently identified through lineage tracing experiments that coupled barcoding and scRNAseq (Rodriguez-Fraticelli et al., 2020) (Figure 4E-F and S4C). Group 1 also displayed a high LSC17 score, a signature derived from functional LSCs isolated from AML patients that predicts poor clinical outcome (Ng et al., 2016) (Figure 4F and S4C). Taken together, these analyses suggest that group 1 cells possess the highest LSC potential within the I1DN progenitor compartment.

We next analyzed the impact of AG-120 on the relative proportion of cells within each group (Figure S5A). As expected from our FACS and morphological observations, AG-120 treatment reduced the fraction of immature cells and conversely increased the abundance of more differentiated cells (Figure S5A-B). Analysis of cell cycle stage inferred from gene expression revealed that AG-120 also greatly increased the percentage of cycling cells, including within the putative LSC group 1 (Figure S5C) (Tirosh et al., 2016). Notably, in both HSCs and LSCs, loss of quiescence is linked with stem cell exhaustion (Lechman et al., 2016; Rodriguez-Fraticelli et al., 2020), suggesting that delayed tumor regression observed in AG120 treated mice is underpinned by a collapse of the malignant hierarchy caused by depletion of functional LSCs.

Many studies have characterized the molecular responses of AML cells to various targeted agents, including IDH inhibitors (Choe et al., 2020; Kats et al., 2017; Shih et al., 2017). What remains unclear however is to what extent the differentiation state of a malignant cell affects its transcriptional response to treatment. To address this question and to understand the effects of AG-120 on different cell types we performed differential gene expression within each progenitor group. This analysis identified partial overlap but also extensive heterogeneity of transcriptional responses, both at the level of individual genes and biological pathways (Figure 5A-C and S5D-F). The transcriptomes of some cell types were more profoundly affected by AG-120 than others, as indicated by the number of DEGs that reached statistical thresholds (Figure S5D). DEGs and signatures related to differentiation that were also evident in bulk RNAseq (Figure 2) were enriched in almost all groups (5C and S5E). In contrast, most progenitor group-specific DEGs that were selectively perturbed only in individual cell types were not detected as differentially expressed in the bulk sequencing data (Figure S5F and Supplemental Table 3). Cell cycle and proliferation related signatures were much more strongly upregulated in group 1 cells; and interferon a and apoptosis signatures were upregulated by AG-120 treatment in some cell types and downregulated in others (Figure 5C). These findings highlight the impact of transcriptional differences between genetically identical leukemic cells on their response to therapy.

**Figure 5:**
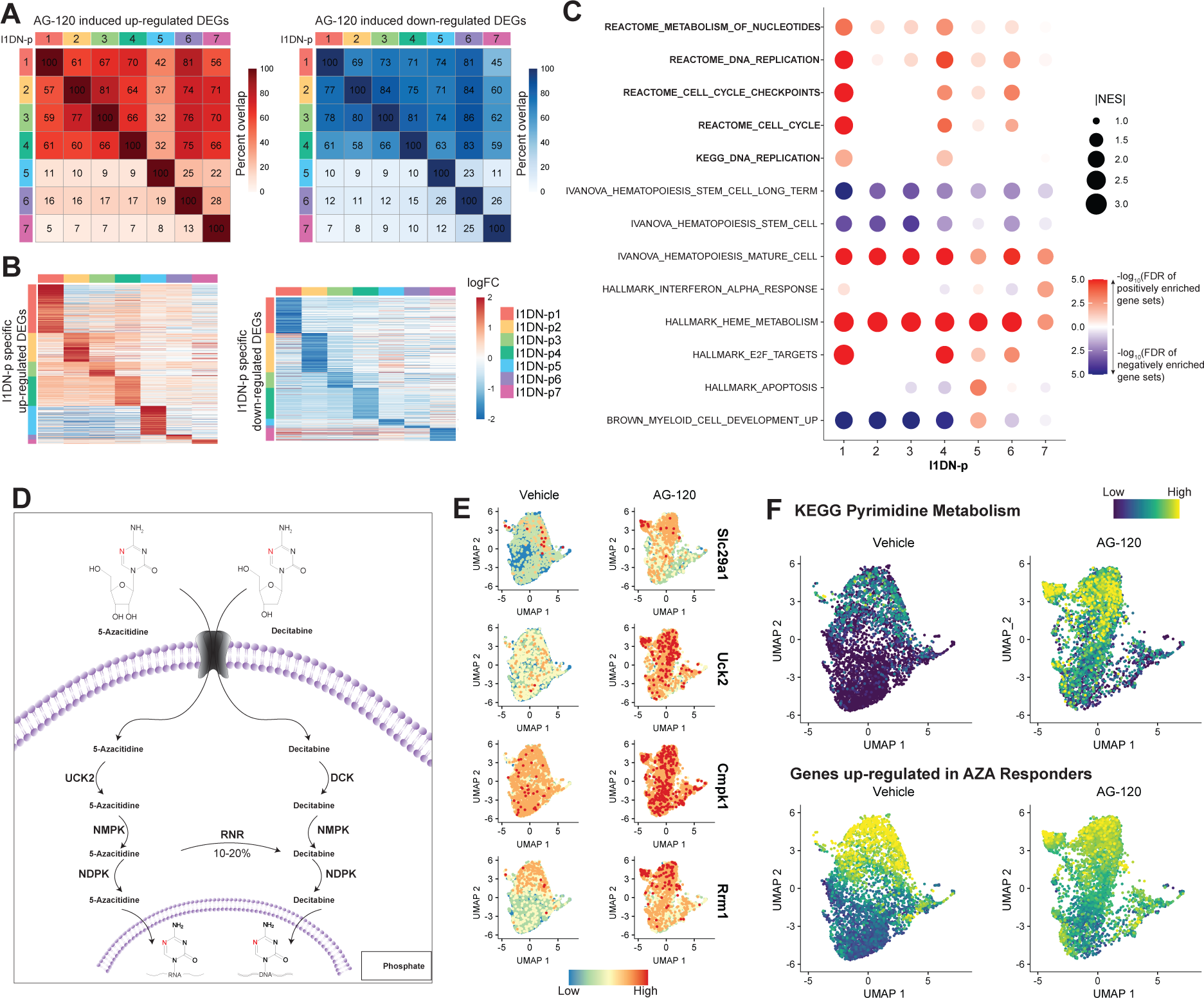
2-HG blockade promotes LSC cycling and upregulation of the pyrimidine salvage pathway. **(A)** Heatmap of the percentage of overlap between the significantly up (left) or downregulated DEGs (right) induced by AG-120 across each I1DN-p group. Percentage overlap was calculated with respect to each column group. **(B)** Heatmap of cluster specific genes that were up (left) or downregulated (right) by AG-120 in different I1DN-p groups. **(C)** Dotplot depicting GSEA signatures enriched upon AG-120 treatment across each I1DNp group. **(D)** Schematic showing the metabolism of hypomethylating agents azacitidine and decitabine. **(E)** UMAP plot of the average expression of specific genes involved in the metabolism of azacitidine. **(F)** UMAP plot of the average expression of the KEGG pyrimidine metabolism signature and genes up-regulated in azacitidine-responder patients (Unnikrishnan et al., 2017).

We then asked whether AG-120 induced transcriptional changes are likely to sensitize AML cells and in particular the LSC compartment to other known therapeutic agents, thereby providing an opportunity for a rationally designed combination strategy. Hypomethylating agents (HMAs) have been proposed as potential combination partners for IDH inhibitors on the basis that the two drug classes promote loss of DNA methylation via distinct pathways (MacBeth et al., 2021). Clinically utilized HMAs such as azacitidine are nucleoside analogues that require DNA replication and enzymes of the pyrimidine salvage pathway for DNA incorporation in order to exert their anti-leukemic effects (Gruber et al., 2020; Gu et al., 2020). In addition to promoting the cycling of quiescent cells, AG-120 also increased expression of pyrimidine metabolism genes that are essential for HMA efficacy including SLC29A1, UCK2 and RRM1 (Figure 5D-F and S5G (Gruber et al., 2020; Gu et al., 2020). An unbiased transcriptional signature of HMA sensitivity derived by comparing HSPCs from myelodysplasia patients that responded or were refractory to azacitidine treatment (Unnikrishnan et al., 2017) was also prominently upregulated in AG-120 treated cells, including the I1DN progenitor group 1 cells (Figure 5F and S5G). Taken together, our data suggests that IDH inhibitors and HMAs are likely to synergize and promote the elimination of quiescent LSCs leading to deep and long-lasting remission.

### AG-120 and azacitidine synergize to provide long-lasting disease control

To test the efficacy of the rationally designed combination of AG-120 and azacitidine we first performed a cross-sectional study in I1DN tumor-bearing mice. We used a modified dosing schedule where AG-120 was administered once per day and azacitidine was administered once per day in cycles of 2 days on and 1 day off. Animals were treated with vehicle, single agents or the combination, and disease progression was assessed on day 10.

Consistent with the reported sensitivity of IDH mutant cancer cells to HMA treatment (MacBeth et al., 2021), azacitidine reduced leukemic burden at levels comparable with AG120. The combination treatment was most effective, reducing overall disease burden and targeting both the leukemic progenitor compartment and mature leukemic cells (Figure 6A-C).

**Figure 6:**
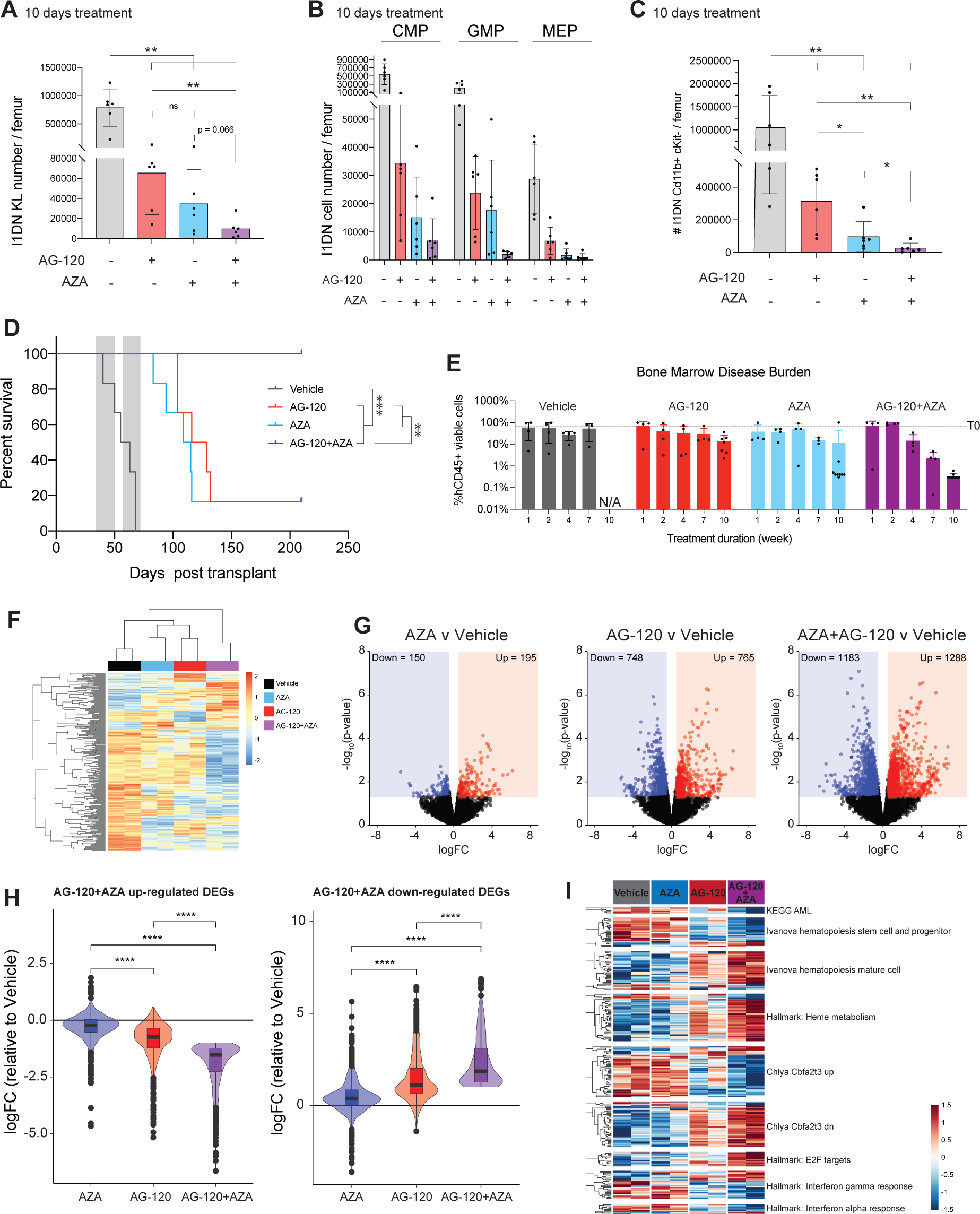
Synergistic activity of AG-120 and azacitidine. **(A-C)** I1DN tumor-bearing mice were treated with AG-120, azacitidine, the combination or vehicle for 10 days and the disease burden in the bone marrow quantified by flow cytometry.

We next tested the impact of the single agent and combination regimens on survival (Figure 6D). Tumor-bearing mice were treated for two 14-day cycles separated by a 7-day recovery period using the modified regimen and monitored for symptom onset and disease progression in the peripheral blood. All mice in the vehicle group reached ethical end point prior to the conclusion of cycle 2. As expected from the cross-sectional study both AG-120 and azacitidine significantly increased survival, although most animals (5/6) in the single treatment arms eventually succumbed to AML. Conversely, all animals in the combination group showed long-term survival with disease control that was sustained for at least 22 weeks after treatment was withdrawn. In all 6 of these mice, FACS analysis of peripheral blood at the end point failed to detect any circulating I1DN cells, potentially suggesting that the animals were leukemia free. To provide orthogonal validation and confirm the relevance of our findings to human AML, we next turned to an *IDH1^R132H^* patient derived xenograft (PDX) AML model that also carried mutations *NPM1^W288fs*12^*, *DNMT3A^A571fs^* and *FLT3^ITD^*. Mice were randomized to treatment cohorts and preassigned groups from each cohort were euthanized at 1, 2, 4, 7 or 10 weeks post enrolment, with the exception of the control group as remaining mice required termination after week 7 due to reaching ethical end point. Disease progression in the bone marrow was quantified by FACS using antibodies directed against human CD45 (hCD45). In this model too, the combination of AG-120 and azacitidine was highly efficacious with all 10 mice analyzed at week 10 having <1% tumor burden (Figure 6E).

We next investigated the consequences of the combined AG-120 and azacitidine treatment on DNA methylation and gene expression using reduced representation bisulfite sequencing (RRBS) and RNAseq. Azacitidine triggers global loss of DNA methylation via degradation of the maintenance DNA methyltransferase DNMT1 that depends on replication and is proportional to drug uptake and incorporation into DNA (Unnikrishnan et al., 2018). IDH inhibitors are also associated with DNA demethylation linked to de-repression of TET2, although the impact of these drugs is less pronounced and more variable, especially in a DNMT3A mutant context (Wang et al., 2021). Moreover, analyses that do not enrich for specific cell types and/or are performed at late timepoints may be confounded by the effects of differentiation.

To capture the acute impact of treatment we analyzed I1DN progenitors isolated from mice that had been treated for five days with vehicle, each single agent alone or the combination (Figure S6A-B). As expected for RRBS we observed greatest sequence coverage in CpG-rich regions (Figure S6C) (Akalin et al., 2012) and thus focused our analysis on comparing the methylation status of promoters. Azacitidine treatment resulted in the fewest differentially methylated promoter regions (FDR < 0.05, 46/13549 promoter DMRs) and had minimal impact on transcription (Figure 6F-G, S6D). Indeed, under azacitidine treatment no genes passed our stringent statistical thresholds (FDR < 0.05) and only 345 DEGs reached significance when the thresholds were relaxed (p < 0.05 and |logFC| > 0.5). AG-120 induced robust gene expression changes (1,513 DEGs at the relaxed threshold) but had only a minor impact on promoter methylation (47/13549 hypomethylated and 23/13549 hypermethylated promoters), suggesting that acute transcriptional perturbation following IDH inhibition in I1DN promoters is not associated with demethylation of gene promoters (Figure 6F-G). Although a lower dose of AG120 was used in this experiment compared with our earlier studies (150mg/kg once daily versus 150mg/kg b.i.d.) gene expression changes were highly concordant (Figure S6E). The combination of AG-120 and azacitidine had the greatest effects on both DNA methylation and gene expression. The majority of differentially methylated promoters were hypomethylated as expected (190/13549 hypomethylated and 58/13549 hypermethylated promoters) (Figure 6FG). Interestingly, dual inhibition of mutant IDH1 and DNMT1 amplified the magnitude of gene expression changes induced by IDH1 inhibition alone (Figure 6H-I, S6F). For example, genes that promote myeloid differentiation were further upregulated, whilst genes that underpin HSC function were further repressed by the combination (Figure 6D). Altogether, our data demonstrate that the AG-120 and azacitidine combination has superior efficacy to either agent alone in aggressive murine and human AML models and is potentially curative in some contexts.

The indicated progenitor (A and B) and mature (C) cell numbers are shown. (n = 6 mice/group; data represented as mean ± SD; for A and C, P-values was calculated using one-tailed, unpaired, non-parametric t-test). **(D)** Kaplan-Meier survival curve of I1DN tumor-bearing mice treated with AG-120, azacitidine, the combination or vehicle. Grey shading denotes treatment. Mice that succumbed to graft vs host disease were censored (n = 6 mice/group; P value was calculated by log-rank test). **(E)** Bone marrow tumor burden (hCD45^+^ cells as percentage of all viable cells) in the mIDH1 AML PDX model treated with AG-120, azacitidine, the combination or vehicle. Mice were culled at pre-defined time-points post enrolment. (n = 4-10 mice/group; data represented as mean ± SD). **(F-I)** RNA-seq and RRBS on cKit+ I1DN cells harvested from the bone marrow of I1DN engrafted mice treated for 5 days. **(F)** Heatmap of significantly methylated promoters (FDR < 0.05) in any drug condition relative to vehicle. **(G)** Volcano plots of DEGs (p < 0.05 and |logFC| > 0.5). **(H)** DEGs induced by the combination of AG-120+AZACITIDINE enhance the effect of AG-120. Violin plots demonstrating that DEGs induced by AG-120+AZACITIDINE are trending in the same direction as the single-agents. P values were calculated using student’s t-test. **(I)** Heatmap of genes from various GSEA pathways that are significant in AG-120+AZACITIDINE. * P < 0.05, **P < 0.01, *** P < 0.001, **** P < 0.0001

## DISCUSSION

Allosteric inhibitors of mutant IDH1/2 hold significant promise for the treatment of AML, however, resistance to single agent therapy develops in almost all patients (Choe et al., 2020). Hence, there is a need to better understand mechanisms that dictate drug response and identify novel combination regimens. While data from clinical trials are accruing, studies in faithful model systems can provide valuable molecular insights that ultimately contribute to optimal drug deployment.

We developed a novel model of IDH1 mutant AML with co-expression of dominantnegative DNMT3A and oncogenic NRAS alleles. AML sequencing studies that have analyzed variant allele frequencies or mutational profiles at single cell resolution have demonstrated that leukemic cells commonly carry 3 or more driver mutations (Miles et al., 2020; Papaemmanuil et al., 2016). Epigenetic mutations such as those in IDH1/2 and DNMT3A frequently co-occur in individual cells and are often present together in dominant leukemic clones suggesting functional co-operation (Miles et al., 2020; Papaemmanuil et al., 2016). NRAS mutations are also common in IDH mutant AML but are usually present in minor sub- clones at diagnosis (Miles et al., 2020; Papaemmanuil et al., 2016). The rapid outgrowth of cells expressing all 3 mutations in primary transplants in our study suggests that they jointly promote leukemia initiation. Our data are in agreement with earlier studies that investigated combinations of epigenetic and signaling mutations (Kats et al., 2017; 2014; Shih et al., 2017; X. Zhang et al., 2016).

The malignant hierarchies in AML differ significantly between patients dictated by various genetic and cell-of-origin factors. In some cases, the majority of leukemic cells resemble early myeloid progenitors, whereas in others the bulk of the tumor is comprised of more differentiated cell types (Miles et al., 2020; Quek et al., 2018; van Galen et al., 2019; Velten et al., 2021). Interestingly, NRAS mutations are associated with a late differentiation block and promote increased CD11b expression in IDH1 mutant AML clones (Miles et al., 2020). Consistent with these observations, we found that I1DN tumors were characterized by a differentiation block at the pre-neutrophil stage, with the majority of leukemic cells expressing CD11b. Thus, the I1ND model recapitulates epigenetic heterogeneity of human IDH/RAS double-mutant AML.

Mice bearing I1DN tumors responded to AG-120 single-agent treatment. Within 5 days of initiating therapy we observed a shift in the leukemic progenitor compartment in the bone marrow to a more differentiated GMP-like immunophenotype. By 4 weeks most of these cells were eliminated, leaving behind only a very small population of residual CMPs. This was accompanied by a progressive increase in differentiated myeloid cells. These observations are consistent with a model whereby IDH inhibitors target multiple levels of the AML hierarchy, limiting the self-renewal of LSCs as well as driving the differentiation of leukemic blasts, leading to disease exhaustion. Our findings underscore the complexity of accurately assessing therapeutic response in the clinical context. Most patients require several months of treatment to achieve their best morphologic response (Amatangelo et al., 2017; DiNardo et al., 2018), likely explained by the time required for LSCs to transit through the blast phase.

The molecular and cellular effects of AG-120 were highly similar to dox-induced genetic depletion of mutant IDH1, formally confirming the on-target activity of the inhibitor. GATA1 was upregulated by AG-120 and implicated as one of the key drivers of the transcriptional response in I1DN progenitors. Mutations in differentiation-associated transcription factors are associated with *de novo* and acquired resistance to IDH inhibitors in patients, suggesting a functional requirement for these pathways in the therapeutic response (Choe et al., 2020; Quek et al., 2018). Mutations in signaling pathways including NRAS are also associated with inferior response to IDH-targeting drugs and are often acquired or enriched at relapse (Choe et al., 2020; Quek et al., 2018). Our data demonstrate that NRAS mutations do not preclude response to single agent IDH inhibition *per se*, although concordant with clinical findings AG-120 failed to completely eliminate the disease.

To explore the transcriptional heterogeneity of leukemic progenitors and its impact on drug response we performed scRNAseq. Recent single-cell lineage tracing studies in HSCs and AML have revealed that cells that are destined for long-term self-renewal or clonal dominance are characterized by an intrinsic and heritable gene expression profile (Fennell et al., 2021; Rodriguez-Fraticelli et al., 2020). Using established signatures from mice and humans we identified a sub-population of quiescent I1DN progenitors that likely represent functional LSCs. The effects of AG-120 on gene expression was distinct in different cell types, suggesting that the maturation status of a leukemic cell is a major factor in dictating its response to IDH inhibition.

An inability to eradicate cancer stem cells and minimal residual disease is a major limitation of many cancer therapies, including targeted agents. The cells that survive initial treatment and ultimately serve as the reservoir for relapse typically represent a minor proportion of the initial tumor and are invisible to methods that analyze bulk cell populations. We found that, AG-120 promoted cycling of LSCs and also increased expression of pyrimidine salvage enzymes in LSCs and other I1DN progenitors, leading us to test the combination of AG-120 with azacitidine. This proved to be highly effective in I1DN AML and in an IDH1 mutant PDX model. Consistent with our hypothesis that AG-120 promotes uptake of azacitidine in immature leukemic cells, the combination treatment caused loss of DNA methylation at gene promoters at a time when single agent treatments had minimal impact.

Notably, synergy between AG-120 and azacitidine has recently been reported in model systems (MacBeth et al., 2021) and patients (DiNardo et al., 2021), although our study is the first to provide a mechanistic basis for LSC targeting.

Together, our findings highlight the importance of understanding and analyzing the impact of epigenetic heterogeneity on drug response. A rapidly growing body of evidence suggests non-genetic mechanisms are a major component of drug resistance in cancer (Lewis and Kats, 2021; Marine et al., 2020). Developing rational strategies that can eliminate cells that have the potential to drive relapse will improve patient outcomes.

## METHODS

### Experimental animals

All animal experiments related to the I1DN mouse model were performed at the Peter MacCallum Cancer Centre and were approved by the Peter MacCallum Cancer Centre Animal Experimentation Ethics Committee. Wild-type C57BL/6, congenic C57BL/6.SJL-Ptprca (referred throughout the text as *Ptprca*) and immune deficient NOD.Cg-Prkdcscid Il2rgtm1Wjl/SzJ (referred to as NSG) mice were purchased from the Animal Resources Centre of Peter MacCallum Cancer Centre or the Walter and Eliza Hall Institute of Medical Research (Melbourne, Australia). All experimental mice were housed at the Peter MacCallum Cancer Centre under specific pathogen-free conditions. Animals were group-housed in individually ventilated micro-isolator cages (6 mice per cage) on a 13-hour light / 11-hour dark cycle. Mice had continuous access to sterilised water and Barastoc irradiated mouse cubes (Ridley AgriProducts). The patient derived xenograft study (PDX) was performed by Champions Oncology (NJ, USA) under approved protocol #1029-018 and commissioned by Agios Pharmaceuticals. NOD.Cg-Prkdcscid Il2rgtm1Sug/JicTac (NOG) female mice were supplied by Taconic and housed in individual HEPA ventilated cages on a 14 hour light / 10-hour dark cycle.

### Retroviral constructs

All retroviruses were based on an MSCV backbone (Clonetech). The doxycycline (doxy)-inducible construct TRI-IDHR132H was generated by replacing the open reading frame encoding MLL-AF9 with an open reading frame encoding the human IDH1R132H allele.

MIT-NRASG12D and MIG-DNMT3AR882H were kind gifts from Johannes Zuber (Research Institute of Molecular Pathology, Vienna, Austria) and Yueying Wang (Shanghai Institute of Hematology, Shanghai, China), respectively.

### Retroviral generation and transduction of murine foetal liver cells

HEK-293T cells, seeded at approximately 50% confluence 24 hrs prior to transfection, were transiently transfected using the calcium phosphate method. A solution of retroviral DNA plasmids (TRI-dsRED-IDH1^R132H^, MSCV-GFP-DNMT3A^R882H^, MSCV-tTA-Nras^G12D^, pclEco packaging vector (Addgene plasmid cat# 12371), 0.25 M CaCl2, 1.25 mM HEPES buffered H2O was added dropwise to an equivalent volume of 2x HEBS buffer (HEPES 50 mM, NaCl 180 mM, Na2HPO4 1.5 mM pH 7) whilst air was bubbled through. The transfection mixture was incubated at room temperature for 20 min then added dropwise to the HEK-293T cells. Embryonic day 13.5-14.5 foetal liver cells (FLCs) were harvested from C57BL/6 embryos and cultured overnight in Dulbecco’s Modified Eagle Medium (DMEM) supplemented with 10% foetal calf serum (FCS), IL-3 (2ng/ml), IL-6 (2ng/ml) and SCF (10ng/ml) prior to transduction. Viral supernatant was collected 48 and 72 hrs post transfection, mixed in a 1:1:1 ratio and spun onto RetroNectin (Takhara)-coated tissue culture plates at 2000g for 1 hr. FLCs were added to the virus-coated plates and centrifuged at 500g for 4 min. Transduced FLCs were then cultured for a further 72 hrs prior to flow-cytometry validation of transduction efficiency and transplant into murine recipients.

### Transplantation and disease monitoring for the I1DN model

To generate murine leukemias, 10^6^ transduced FLCs (consisting of a mixture of untransduced, single-, double- and triple-transduced cells) were transplanted into sublethally irradiated (6.5Gy administered as a single dose) *Ptprca* recipients by intravenous injection into the tail vein. For serial transplantation experiments, bone marrow or spleen cells from moribund animals were collected and cryopreserved in FCS supplemented with 10% DMSO. Cryopreserved cells were thawed and washed in Dulbecco’s phosphate-buffered saline (PBS), and 1-2 x 10^5^ viable cells were transplanted into sub-lethally irradiated (6.5Gy administered as a single dose) *Ptprca* recipients, or unirradiated NSG recipients, by intravenous injection into the tail vein. To reduce the occurrence of graft-vs-host disease (GvHD), I1DN cells were stained for 15 min on ice with a lymphoid lineage cocktail containing antibodies against Tcrb (eBioscience 47-5961-82), B220 (Invitrogen 47-0452-82) and NK1.1 (BD Pharmingen 560618), then washed and resuspended in FACS buffer with DAPI. Cells were sorted on the BD FACSAria Fusion 5 for the dsRED+/GFP+/DAPI-/lymphoid lineage- population before being washed, resuspended in PBS and transplanted into NSG mice.

Animals were closely monitored for clinical signs of disease development (weight loss, lethargy, hunched posture) by animal house technical staff who were blinded as to the conditions of the experiment. Peripheral blood was routinely collected by tail vein incision into EDTA coated Microvetteâ capillary blood collection tubes (Sarstedt AG & Co) to assess changes in blood cell counts and detect presence of leukemic cells by flow cytometry. Animals were euthanized at ethical end point based on clinical symptoms and overall survival rates were assessed using Kaplan-Meier analysis.

### Limiting dilution transplant assay

I1DN cells were stained for 15 min on ice with antibodies against cKit (BD Pharmingen 553356), CD11b (Thermo Fisher 47-0112-82), CD34 (BD Bioscience 562608), FcgR-III (BD Optibuild 740851) and Sca1 (BD Pharmingen 558162), then washed and resuspended in FACS buffer. Cells were sorted on the BD FACSAria Fusion 5 for I1DN CMP (GFP+/CD11b/CD117+/Sca1-/CD34^mid^/ FcgR-III^mid^) and GMP (GFP+/CD11b-/CD117+/Sca1-/CD34^high^/ FcgR-III^high^) cells. Sorted cells were washed and resuspended in PBS at 4x10^5^/mL, 4x10^4^/mL, 4x10^3^/mL. 200 µL of cells were transplanted into NSG mice (N = 3 / cell type / cell number).

### I1DN therapy experiments

Mice assigned to doxycycline (DOX) treated cohorts were provided with DOX supplemented chow (600 mg/kg) (Speciality Feeds) and DOX water (DOX 0.2% w/v, sucrose 2% w/v) for the duration of the experiment. AG-120 (150 mg/kg) was administered by twice daily oral gavage of an amorphous solid dispersion with hypromellose acetate succinate (HPMCAS) as the carrier for the active pharmaceutical ingredient. AG-120 was prepared by resuspending 3% (w/v) AG-120-SDD, 0.5% (w/v) methylcellulose and 0.2% (w/v) Tween80 in purified water. AG-120-SDD (Agios Pharmaceuticals) contained the active pharmaceutical ingredient (AG-120) in a 1:1 (w/w) ratio with HPMCAS. Vehicle was comprised of 1.5% (w/v) HPMCAS, 0.5% (w/v) methylcellulose and 0.2% (w/v) Tween80 in purified water. Vehicle and AG-120 were administered to recipient mice by oral gavage twice daily at doses relative to body weight (10 mL/kg) for the duration of the experiment defined. The treatment protocol was modified for the Azacitidine and AG-120 combination study to comply with ethics requirements. Azacitidine was prepared fresh daily in PBS to a final concentration of 0.1 mg/mL and was administered by intraperitoneal injections according to body weight (1 mg/kg). Azacitidine treatment was administered once per day, in cycles of 2 days on and 1 day off, for 14 days. Mice received a 1-week treatment break, followed by another 14 days of the same azacitidine cycle. During each treatment period, AG-120 was administered once every day (150 mg/kg). On days where the combination treated group received both azacitidine and AG-120, drugs were administered at least 6 hrs apart.

### PDX therapy experiments

Within 4 hrs prior to inoculation, female NOG mice were sub-lethally irradiated with 1.5Gy whole body irradiation. AML cells #CTG-2227 carrying *IDH1^R132H^*, *NPM1^W288fs*12^*, *DNMT3A^A571fs^* and *FLT3^ITD^* mutations were thawed at 37°C and diluted to a concentration of 10 x 10^6^ cells/mL.UCHT1 (BioXCell) will be diluted to 1 mg/mL solutions in sterile PBS containing 2% FBS and stored at 4°C until use. UCHT1 antibody was incubated with cells in suspension on ice for 30 minutes at a concentration of 1 μL/10^6^ and 2 x 10^6^ cells were transplanted into each recipient via injection into the lateral tail vein. Animals were treated with azacitidine at a dose level of 3mg/kg with a volume of 10mL/kg IP/Q3D, AG-120 at a dose level of 900mg/kg with a dose volume of 10mL/kg PO/QD or the combination.

### AG-120 and 2-HG measurements in mouse serum

50 µL of peripheral blood was centrifuged at 2000g for 4 min at 4 °C to separate the plasma from the cells. 20 µL of plasma supernatant was carefully transferred into a new eppendorf tube and immediately stored at -80 °C. Bone marrow from a single femur was isolated by pulse centrifugation into an eppendorf tube and stored at -80 °C. The concentrations of AG-120 in plasma and tissue samples were determined using non-validated liquid chromatography with tandem mass spectrometry (LC-MS/MS) methods. Tissue samples were homogenized using a FastPrep homogenizer for 60 seconds, with 10 volumes (volume- byweight [v/w]) of methanol:water (80:20 [v/v]) to get a homogenate with a dilution factor of 11. Calibration standards and quality control (QC) samples were prepared in blank mouse plasma. A 10-µL aliquot of calibration standards, QCs, unknown plasma and tissue homogenate were mixed with 200 µL of acetonitrile containing the internal standard (IS) AGI- 0018070 (25 ng/mL) for protein precipitation. The mixture was vortexed at and centrifuged. A 100-µL aliquot of supernatant was mixed with 100 µL of water with 0.1% formic acid and vortexed to mix and was analyzed on a SCIEX Triple Quad™ 6500+ with Exion LC™ AD system. A reversed-phase gradient method using a Waters ACQUITY UPLC HSS T3 Column (100Ã, 1.8µm, 2.1 mm X 50 mm) maintained at 50 °C, provided chromatographic separation. Water with 0.1% formic acid and acetonitrile with 0.1% formic acid were used as mobile phase A and B respectively, at a total flow rate of 600 μL/min. AG-120 and the IS were ionized under a positive ion electrospray mode and detected through the multiple-reaction monitoring (MRM) transitions of m/z 583.2/214.0 and m/z 587.3/214.0. Data was acquired using Analyst 1.6.3 (AB Sciex, Foster City, CA). The standard curve had a coefficient of determination (R2) value >0.98 in a linear regression with 1/X2 weighting. The quality control samples had a precision and accuracy within 20% of theoretical values. The peak area ratios of analyte relative to internal standard were used for AG-120 quantitation. Linearity was achieved in the AG-120 concentration range from 1 ng/mL to 30,000 ng/mL.

### Flow cytometry analysis and cell sorting

A 10 mL single cell suspension of bone marrow in FACS buffer (2% heat inactivated FCS in PBS) from a single intact femur per mouse was obtained by mechanical dissociation, with a mortar and pestle, which was subsequently filtered through a 70 µm cell-strainer (Greiner Bio-One). One eighth of the single cell suspension obtained from a single femur per mouse was isolated for quantitative flow-cytometry. Splenocytes were isolated by pressing the spleen through a 70 µm cell-strainer with a volume of FACS buffer relative to spleen weight (1 mg / 40 µL). 100 µL (equivalent to 2.5 mg spleen) of the splenocyte cell suspension was used for quantitative flow-cytometry. For flow cytometric analysis and cell sorting, single-cell suspensions of whole blood, bone marrow and spleen were incubated in ACK red cell lysis buffer (150 mM NH4Cl, 10 mM KHCO3, 0.1 mM EDTA) for 2 min and then washed in FACS buffer (PBS, 2% FBS). Cells were then re-suspended in FACS buffer (2% heat inactivated FCS in PBS) and stained with fluorophore-conjugated antibodies targeted against cell surface markers. All antibodies in Table 1 were diluted 1/200, except for CD34 which was diluted 1/100 and Ly-6G which was diluted 1/400. Cells were stained for 15 min on ice or 5 min in a 37°C water bath, washed then resuspended in FACS buffer. Cells were analysed on the BD LSR Fortessa X-20 or BD FACSymphony or sorted on the BD FACSAria Fusion 5 or BD FACSAria Fusion 3 (BD Biosciences). Data was analysed using FlowJo (v 10.4).

### Western blot

Bone marrow and spleen derived I1DN cells sorted by FACS, were pelleted by centrifugation (450 x g, 4 min, 4 °C). Pelleted cells were washed with PBS and lysed for 1 hr on ice in RIPA buffer (NaCl 150 mM, MgCl2 2 mM, Triton X-100 1% (v/v), Sodium Deoxycholate 0.5% (w/v), SDS 0.1% (w/v), Tris-Cl 50 mM pH 8.0) with 1 x protease inhibitor cocktail (Roche Diagnostics) and Bezonase 1.25 U/uL (Merk Millipore). Protein concentration was quantified using the Microplate Procedure with the Pierce BCA Protein Assay Kit (Thermo Scientific) according to the manufacturer’s instructions and used to calculate the concentration of the samples. 20 – 40 μg of protein lysates were diluted to 1 – 2 μg/μL with PBS and 6 x SDS (Tris base 375 nM pH 6.8, SDS 12.3% (w/v), Glycerol 25.8% (v/v), bromophenol blue 600 μg/mL, 2-mercaptoethanol 60 μL/mL). Samples were incubated at 95 °C for 5 minutes prior to being resolved on a 4 – 15% Mini-PROTEAN TGX Precast Protein Gels (Bio-rad) and transferred onto a methanol activated Immobilon-P PVDF membrane (Millipore) by electroblotting in transfer buffer (Tris base 25 mM, glycine 1.92 M, methanol 10% (v/v)) for 60 minutes at 4 °C. Membranes were blocked in 5% (w/v) skim milk powder in 1 x TBS/T (Tris base 50 mM pH 7.6, NaCl 150 nM, Tween20 0.1% (v/v)) for 1 hour at room temperature prior to incubation in primary antibody (Table 2) diluted in 5% (w/v) skim milk powder in 1 x TBS/T overnight at 4 °C. Membranes were washed with 1 x TBS/T for 15 min in 5 min intervals before incubation in HRP-conjugated secondary antibody diluted in 5% (w/v) skim milk powder in 1 x TBS/T for 1 hour at room temperature. Membranes were subsequently washed with 1 x TBS/T for 30 min in 10 min intervals prior to incubation in ECL^TM^ (GE Healthcare, cat# RPN2106) and exposure by film (FUJIFILM Super RX) using an AGFA CP1000 developer (AGFA).

### RNA sequencing and analysis

1 – 5 x 10^5^ cells (GFP^+^cKit^+^CD11b^-^) per mouse were isolated by FACS, washed with PBS and pelleted by centrifugation (450 x g, 10 min, 4 °C) prior to being stored at -80 °C in TRIzol Reagent (Invitrogen). RNA was extracted with the Direct-zol RNA Microprep kit as per manufacturer’s instructions (Zymo Research R2062). Purified RNA was quantified using the TapeStation 2200 system (Aligent) with RNA HS screentape (Aligent) according to the manufacturer’s instructions. RNA was stored at -80 °C until required. The Molecular Genomics Core of the Peter MacCallum Cancer Centre used the QuantSeq 3’ mRNA-seq Library Prep Kit for Illumina (Lexogen 015), according to the manufacturer’s instructions, to generate the sequencing libraries from the purified RNA. 75 bp single end reads with a depth of 6 – 10 million reads per sample were generated using the Illumina NextSeq500.

Sequencing reads were demultiplexed using bcl2fastq (v 2.17.1.14), low quality (Q < 30) reads removed, and trimmed at the 5’ and 3’ ends using cutadapt (v 1.14) (Martin, 2011) to remove adapter sequences and poly-A-tail derived reads respectively. Sequencing reads were mapped to the mouse reference genome (mm10) using HISAT2 (v 2.1.0) (Kim et al., 2015) and counted using the featureCounts command of the Subread package (v 1.6.3) (Liao et al., 2014). Read normalisation and differential gene expression analysis was performed in R (v 3.6.2) using R packages limma (v 3.42.2) (Ritchie et al., 2015) and EdgeR (v 3.28.1) (McCarthy et al., 2012). RCisTarget (v 1.11.10) (Aibar et al., 2017) was used for motif enrichment analysis. Barcodeplots (limma) were used to visualize gene set enrichment and rotating gene set testing (using the “fry” function from the limma package) was used to test for gene set enrichment. R packages pheatmap (v 1.0.12), ggplot2 (v 3.2.1), ggrepel (v 0.8.1), RColorBrewer (v 1.1-2) and limma (v 3.42.2) (Ritchie et al., 2015) were used to generate figures.

### RRBS sequencing and analysis

5 x 10^4^ cells (GFP^+^cKit^+^CD11b^-^) per mouse were isolated by FACS, washed with PBS and pelleted by centrifugation (450 x g, 10 min, 4 °C). DNA was extracted using the QuikDNA Microprep Kit (Zymo Research D3020) as per the manufacturer’s instructions. Diagenode’s Premium RRBS kit (C02030033) was used according to the manufacturer’s instructions to generate the RRBS libraries from the purified DNA. Each sample contained methylated and unmethylated spike-in controls, as per the instructions in Diagenode’s Premium RRBS kit. Libraries were sequenced on the Illumina Novaseq6000 with 150 bp paired-end reads and a depth of at least 30 million reads per sample.

Sequencing reads were demultiplexed using bcl2fastq (v 2.17.1.14), low quality reads removed, and trimmed with TrimGalore (v 0.4.4) using --paired and --rrbs settings. Bismark (v 0.22.3) (Krueger and Andrews, 2011) was used to map the spike-in controls and samples using default parameters, and samtools (v 1.4.1) (Li et al., 2009) was used to sort and index bam files. Bismark (v 0.22.3) (Krueger and Andrews, 2011) was used to extract coverage files, and differential analysis was performed in R (v 4.0.2) using the standard edgeR (v 3.32) workflow for differential methylation analysis (Chen et al., 2017). Briefly, CpGs were annotated to the closest TSS using edgeR’s nearestTSS function and subset to include on CpGs that lie within 2kb upstream and 1kb downstream of any given TSS, then counts were summed over each gene’s TSS, promoter regions with low coverage removed (promoters retained when methylated+unmethylated counts ³ 10 in every sample), and dispersion estimation and model fitting as per edgeR guidelines.

### Exome sequencing and analysis

Sequencing reads were aligned using BWA-MEM(Li, 2013) using reference genome GRCm38.73, and the aligned reads were processed through three variant callers: MuTect v1.17 (Cibulskis et al., 2013), Vardict v1.4.6 (Lai et al., 2016), and GATK-Mutect2 v3.9.0 (Van der Auwera et al., 2013). Resulting variants were annotated using Variant Effect Predictor (VEP) release 90 (McLaren et al., 2016). Other utility software involved in the pipeline includes BEDTools v2.21.0(Quinlan and Hall, 2010) for region filtering, BCFtools v1.3.1(Danecek et al., 2021) for variant processing and filtering, and Tabix v0.2.5(Li, 2011) for indexing. In R (v 4.0.2), variants detected in at least 2 of the variant callers were retained, low quality (QUAL < 30) and low read depth variants (PMCDP < 10) were excluded and variants with VAF < 0.3 in the I1DN TE leukemias were excluded. Additionally, for variants that affected multiple transcripts of the same gene, the variant predicted to have the greatest impact was retained and the rest were excluded. ggplot2 (3.3.3) was used for figure generation.

### Single-cell RNA sequencing and analysis

I1DN progenitor cells were FACS sorted into PBS/2%FBS. Cells were washed once with RNase free PBS/1%BSA and resuspended in PBS/1%BSA+0.2U/ml RNase inhibitor (Protector RNAse inhibitor, Sigma) at a final concentration of 1000 cells/µL. Cell suspensions were loaded onto separate channels of the 10XChromium Single Cell Chip and single cells were captured in droplet emulsions using the 10X Chromium Single-Cell Instrument. Reverse transcription, cDNA amplification and library preparation were performed using the Chromium Single Cell 5′ Library & Gel Bead Kit (10X Genomics PN-1000006). Libraries were pooled and sequenced on the Illumina NovaSeq 6000 system with 150 bp paired end reads to a depth of ∼50,000 reads per cell.

Count matrices were generated from demultiplexed scRNA-seq fastq files using the 10X Genomics Cell Ranger v3.1.0 count pipeline against the mm10/GRCm38 genome. scRNA-seq quality control, normalization, data integration, dimensional reduction, k-nearest neighbor graph construction and clustering were performed using Seurat (v 4.0.4) (Butler et al., 2018) in R (v 4.0.2). Low quality cells were removed by filtering out cells that had fewer than 1000 genes, fewer than 2000 unique molecular identifiers (UMIs), or greater than 15% mitochondrial RNA content. Seurat’s CellCycleScoring function was used to assign cell cycle phase using Seurat’s inbuilt list of cell cycle genes(Butler et al., 2018; Tirosh et al., 2016). Seurat’s SCTransform function was used for data normalization with percentage mitochondrial reads, and differences in cell cycle phase amongst proliferating cells (S and G2M cell cycle phase scores) included in the regression model as sources of technical variation to remove. Datasets were integrated using canonical correlation analysis (CCA) within Seurat using default parameters. The first 40 principal components were used to compute a non-linear dimensional reduction using the Uniform Manifold Approximation and Projection (UMAP) method, and clustering was performed at a resolution of 0.4. Clusters with fewer than 100 cells were combined with larger clusters based on correlation analysis of the cluster-average expression of the top 500 most variably expressed genes. Pseudo-bulk RNAseq was performed for differential gene expression analysis between treatment conditions. Counts were summed across I1DN progenitor groups for each treatment condition. Differential gene expression within each I1DN progenitor group was performed with edgeR (v 3.32.1) (McCarthy et al., 2012). Gene ontology and GSEA analyses were performed with R packages clusterProfiler (v 3.18.0) (Yu et al., 2012), and fgsea (v 1.16.0) (Korotkevich et al., 2021), respectively. Seurat, ggplot2 (3.3.3) and pheatmap (1.0.12) were used for figure generation.

## DATA AVAILABILITY

Processed and unprocessed data for RNAseq, scRNAseq and RRBS will be deposited to GEO and accession numbers provided prior to publication. All other data is available from the corresponding author on request.

## COMPETING INTERESTS

LMK has received research funding and/or consultancy payments from Agios Pharmaceuticals, Celgene Corporation and Servier Pharmaceuticals. JS has received research funding from BMS/Celgene, Amgen and Astex Pharmaceuticals Inc; and served on the advisory boards of Astellas, Novartis and Mundipharma.

## ACKNOWLEDGEMENTS

We thank members of the molecular genomics, animal and flow cytometry core facilities at the Peter MacCallum Cancer Centre for technical support, Brian Liddicoat for assistance with bisulfite sequencing and Dylan Marchione and Emma Dion for critical review of the manuscript. This work was supported by a research grant from the National Health and Medical Research Council of Australia (APP1099160). The following authors were supported by fellowships: LMK from the Victorian Cancer Agency (MCRF15003), ACL from the Brazis family, JS from the Medical Research Future Fund of Australia (GNT1160133), LGM from The Lorenzo and Pamela Galli Medical Research Trust. EG was supported by the Australian Postgraduate Award.

## AUTHOR CONTRIBUTIONS

Conceptualization and design: EG, RWJ and LMK. Experiments and data analysis: EG, JS, ACL, RF, RC, LGM, AJR, EV, PF, KS, LJ, AT, NT, JL and LMK. Technical expertise and essential reagents: BN, SD, AT and MH. Supervision: JS, RWJ and LMK. Writing and editing: EG, RWJ and LMK.

