## Supplemental files for "Inhibition of mutant IDH1 promotes cycling of acute myeloid leukemia stem cells"

### Supplementary Figure 1

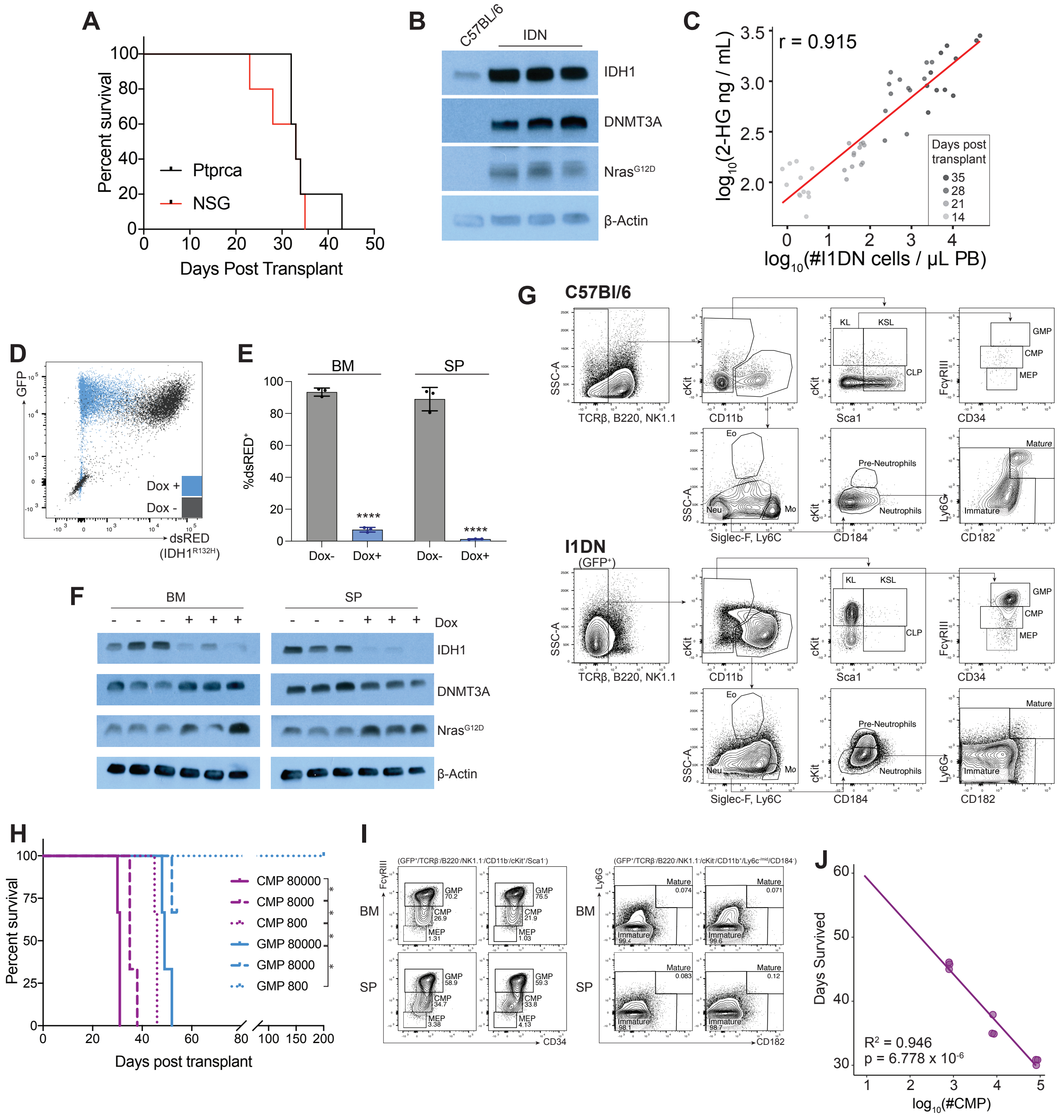


**Figure S1A: Development and characterization of the I1DN AML model.**

**(A)** Kaplan-Meier curve of syngeneic (*Ptprca*) and immune compromised (NSG) recipient mice transplanted with I1DN tumors (n = 6 mice/group). **(B)** Western blot demonstrating elevated expression of IDH1, DNMT3A and NRAS in the bone marrow of I1DN transplanted recipients compared with control C57BL/6 bone marrow. **(C)** Relationship of plasma 2-HG concentration and disease burden in I1DN tumor-bearing mice. **(D)** Representative FACS plots showing immunophenotype of I1DN leukemic cells in the spleen compared with control spleen from a non-leukemic mouse **(E)** Significant reduction in the proportion of leukaemic (GFP+) cells expressing dsRED isolated from IDN transplanted recipients following 5 days of DOXsupplementation. N = 3/group. P value calculated using two-tailed, unpaired, non-parametric t-test (**** $P < 0.0001$). **(F)** western blotting of sorted GFP+ cells harvested from the bone marrow or spleen of leukemic mice untreated or treated with dox for 5 days. **(G)** Representative FACS plots showing immunophenotype of I1DN leukemic cells in the spleen compared with control spleen from a non-leukemic mouse. **(H)** Kaplan-Meier survival curve of NSG recipient mice transplanted with the indicated number of FACS isolated I1DN CMP or I1DN GMP cells (n = 3 recipients/number of CPMs or GMPs transplanted; 2 recipients in the group transplanted with 8000 GMPs were euthanized due to fighting and censored from the graph). **(I)** Representative FACS plots of the bone marrow and spleen demonstrating that the entire leukemic hierarchy is recapitulated following transplant of I1DN CMPs. **(J)** Inverse relationship between survival and number of sorted I1DN CMP cells transplanted (n = 3 recipients/number of CPMs transplanted). BM – bone marrow; SP – spleen; dox – doxycycline.

### Supplementary Figure 2

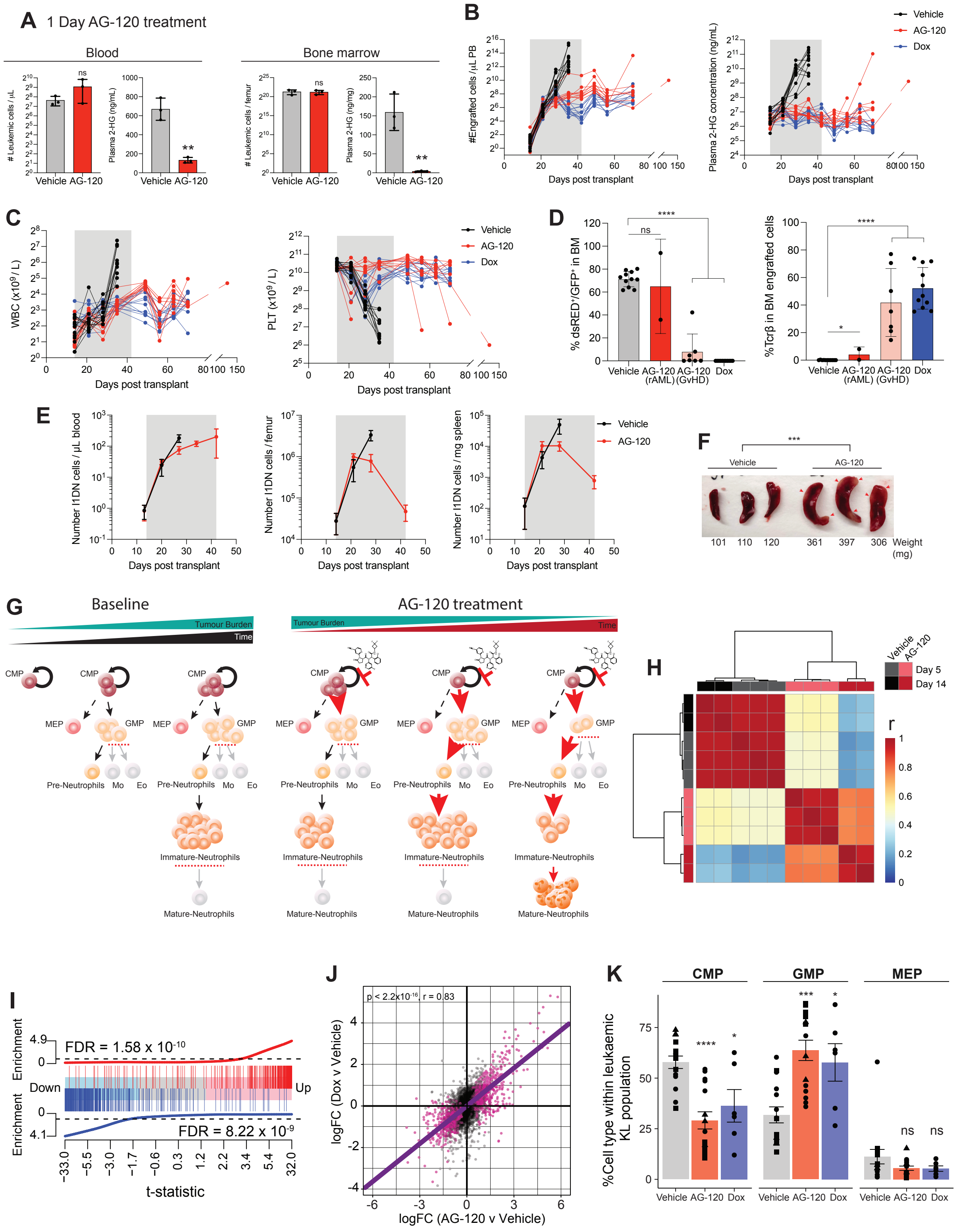

**Figure S2A: Inhibition of mutant IDH1 drives exhaustion of the leukemic hierarchy.**

**(A)** 2-HG concentration in the plasma (left panel) or bone marrow (right panel) of I1DN tumor bearing mice treated for 1 day with AG-120 or vehicle. Total number of leukemic cells were comparable between the AG-120 and vehicle treated samples (n = 3 mice/group; data represented as mean ± SD; P values calculated using two-tailed, unpaired, non-parametric student's t-test). **(B)** Number of engrafted cells quantified by FACS and plasma 2-HG concentration in the peripheral blood of individual I1DN tumor bearing mice treated with AG120, doxycycline or vehicle. Grey shading denotes treatment (n = 11-12 mice/group). **(C)** Number of white blood cells and platelets in the peripheral blood of individual I1DN tumor bearing mice treated with AG-120, doxycycline or vehicle. Grey shading denotes treatment (n = 11-12 mice/group). **(D)** I1DN tumor-bearing mice culled at ethical end point. Mice that succumbed to graft-vs-host-disease had a comparably low percentage of I1DN cells in the bone marrow and an increased number of cells expressing the T cell marker TCRb (data represented as mean ± SD). See also Table S1. P values calculated using two-tailed, unpaired, nonparametric t-test. **(E)** Quantification of I1DN cells in the bone marrow, spleen and peripheral blood of mice treated with AG-120 or vehicle for the indicated period (n = 3-4 mice/group/timepoint; data represented as mean ± SD). **(F)** Photographs of spleens from I1DN leukemic mice treated with AG-120 or vehicle for 5 days. Arrows indicate macroscopic blast infiltrate. **(G)** Schematic depicting the effects of IDH1 inhibition of the AML hierarchy. **(H-J)** RNAseq analysis of sorted I1DN progenitor cells. **(H)** Heatmap of the pairwise Pearson's correlation coefficients of the top 1000 most variably expressed genes. Each column/row represents tumors harvested from individual mice, except for day-14 AG-120-treated samples which encompass cells pooled from two 2 mice due to low yield. **(I)** Barcode plot demonstrating enrichment of signatures derived from mice bearing mutant IDH2 leukemias treated with the IDH2 inhibitor AG-221 (Kats et al., 2017) in mice engrafted with I1DN leukemias and treated with AG-120. **(J)** I1DN leukemic mice were treated with AG-120, dox (genetic de-induction of $IDH1^{R132H}$) or vehicle for 5 days. FACS analysis showing differentiation within the leukemic progenitor compartment (data collected from 3 separate experiments, depicted by different shapes; data represented as mean ± SD). **(K)** Correlation plot of the DEGs induced by 5 days of IDH1$^{R132H}$ inhibition (AG-120) or genetic de-induction (DOX) in the CD117+ leukaemic cells isolated from the bone marrow. N = 2 recipients / group. Purple circles represent significant DEGs (p-value < 0.05) in either comparison. Line of best fit calculated for significant DEGs in either comparison. Error bars represent mean ± SD.

BM – bone marrow; PB – peripheral blood; WBC – white blood cells; PLT – platelets; dox – doxycycline; CMP – common myeloid progenitor; GMP – granulocyte-macrophage progenitor; MEP – megakaryocyte-erythroid progenitor. * p < 0.05, ** p < 0.01, *** p < 0.001, **** p < 0.0001.

### Supplementary Figure 3

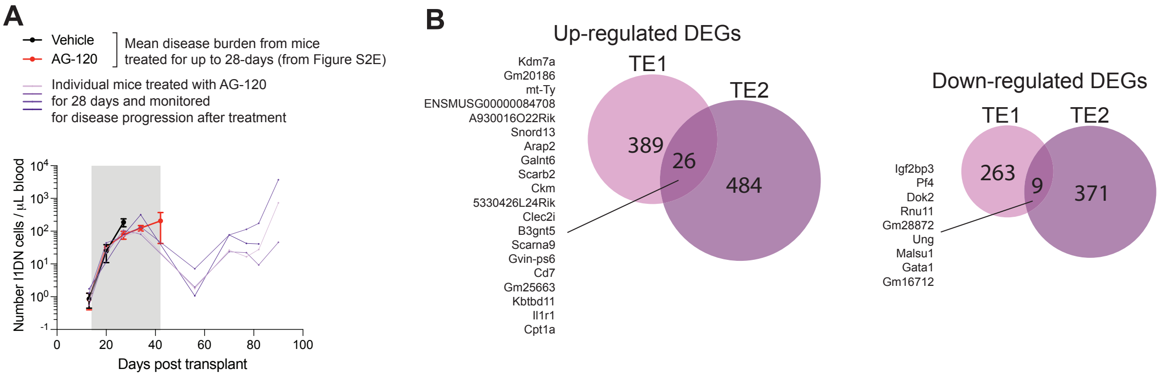


**C**

**TE1 up-regulated DEGs**

| Rank | Motif | Transcription factor | NES |
|---|---|---|---|
| 1 | 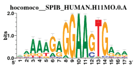 | Spib (inferredBy_Orthology) | 6.13 |
| 2 | 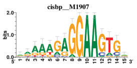 | Spi1 (directAnnotation). | 6.13 |
| 3 | 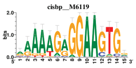 | Spi1 (inferredBy_Orthology). | 6.06 |

**TE2 up-regulated DEGs**

| Rank | Motif | Transcription factor | NES |
|---|---|---|---|
| 1 | 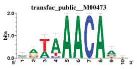 | Foxo1 (directAnnotation). | 5.68 |
| 2 | 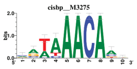 | Foxo1 (inferredBy_Orthology). | 5.68 |
| 3 | 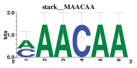 | Zfp128 (inferredBy_Orthology). | 5.6 |

**Figure S3: Characterization of AG-120 resistant I1DN tumors.**

**(A)** Quantification of I1DN cells in the peripheral blood of mice treated with AG-120 (red) or vehicle (black) for the indicated period (n = 3-4 mice/group/timepoint; data represented as mean ± SD). Purple lines indicate peripheral blood tumor burden of 4 individual AG-120 treated mice that received the full 28-day therapy and were monitored for relapse. **(B)** Venn diagram of overlapping DEGs. **(C)** Enrichment of transcription factor binding sites identified within the proximal promoters of the TE up-regulated DEGs identified by RcisTarget. Ranking was determined by normalized enrichment score (NES).

### Supplementary Figure 4

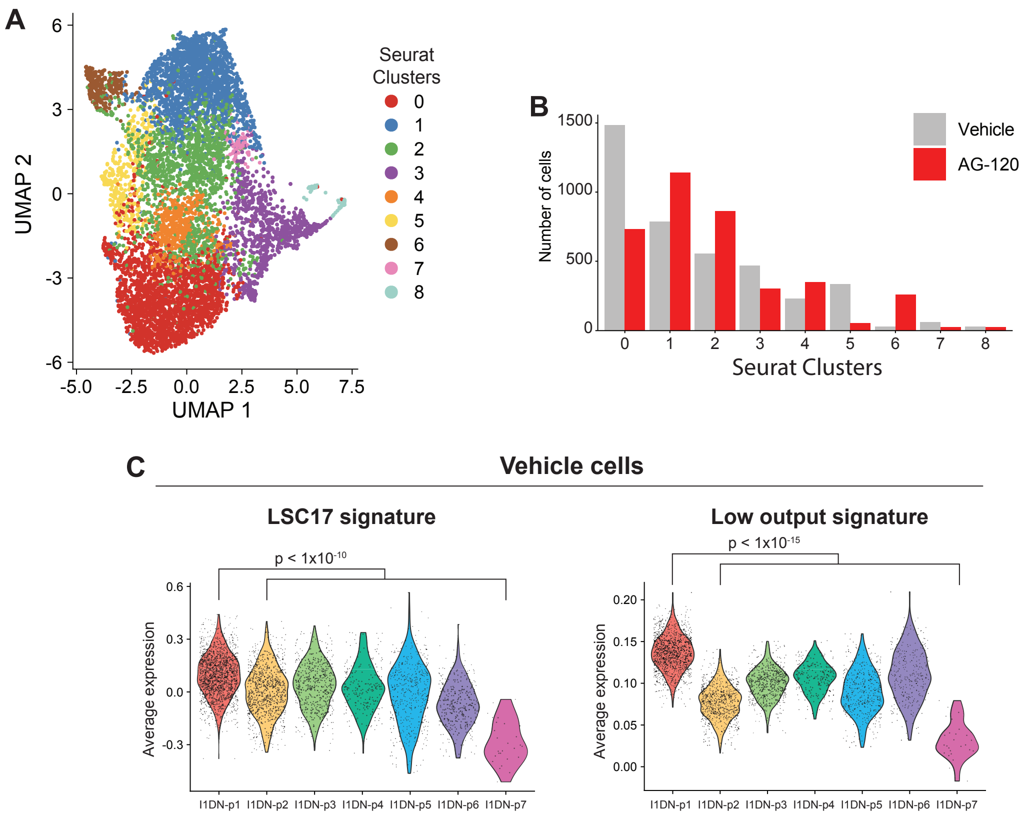


**Figure S4: Transcriptional heterogeneity of I1DN progenitor cells.**

**(A)** I1DN progenitor cells from mice treated with AG-120 or vehicle for 5 days were analyzed by scRNAseq. UMAP plot of integrated datasets labelled by Seurat clusters at a resolution of 0.4. **(B)** Number of cells within each Seurat cluster by treatment condition. **(C)** Violin plot of average expression of hscScore (Hamey and Göttgens, 2019), "low-output" HSC signature (Rodriguez-Fraticelli et al., 2020) and LSC17 score (Ng et al., 2016).

### Supplementary Figure 5

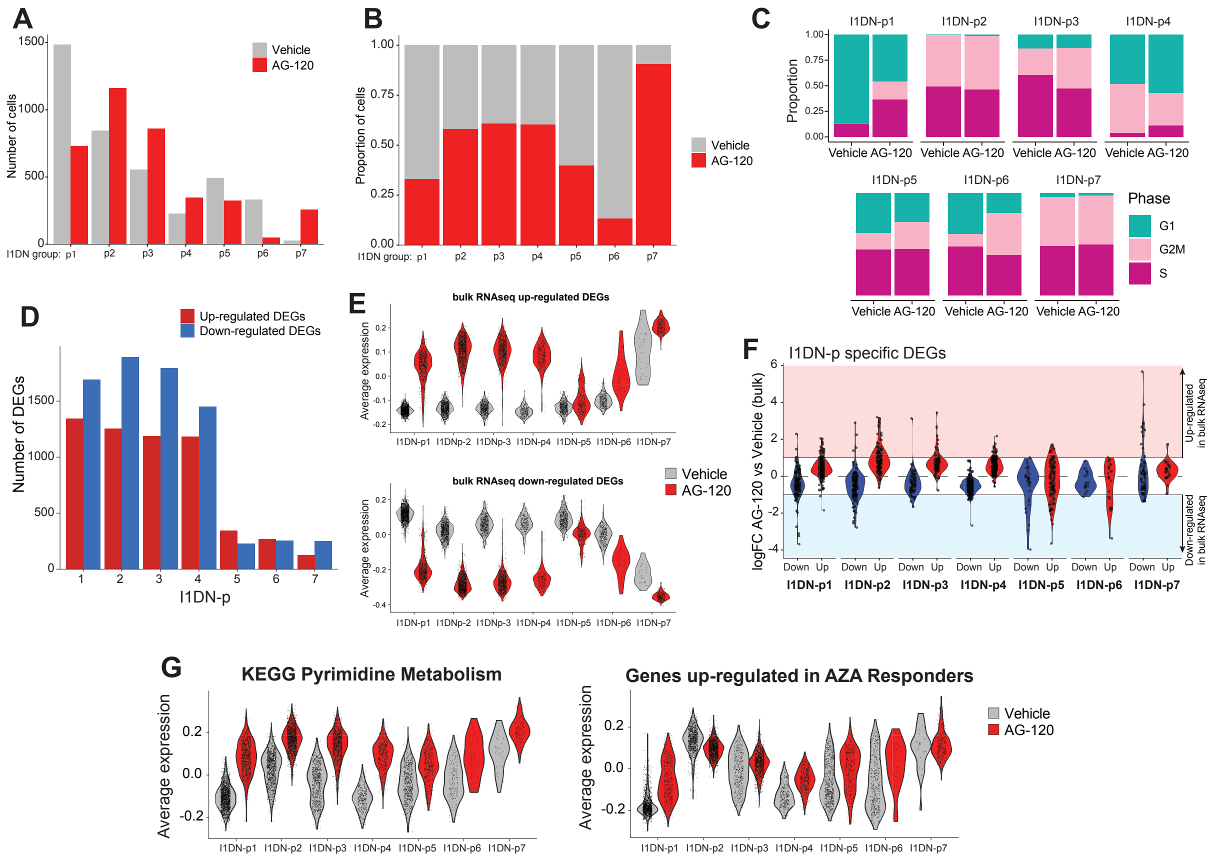


**Figure S5: Heterogeneity of molecular responses in I1DN progenitor cells.**

**(A-B)** Number **(A)** and proportion **(B)** of cells from each treatment condition by progenitor group. **(C)** Proportion of cells within each phase at of the cell cycle in each progenitor group by treatment condition. **(D)** Number of AG-120-induced DEGs within each progenitor group. **(E)** Violin plots of the average expression of DEGs identified in the bulk RNAseq across each progenitor group. **(F)** Violin plots of the bulk RNAseq logFC for genes identified as I1DN progenitor group-specific AG-120-induced DEGs. **(G)** Violin plots of the average expression of genes in the KEGG pyrimidine metabolism signature (left) and genes up-regulated in AZACITIDINE-responder patients (Unnikrishnan et al., 2017) (right).

### Supplementary Figure 6

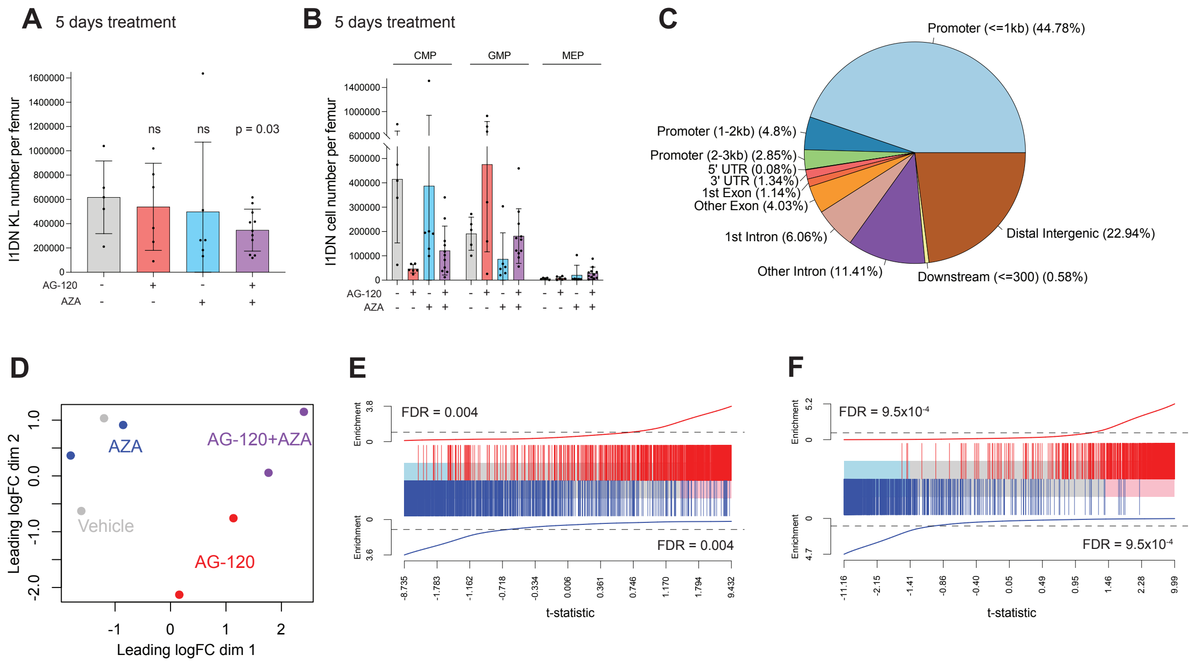


**Figure S6: Synergistic activity of AG-120 and azacitidine in AML.**

**(A-B)** Number of I1DN cKit+ cells following 5 days of treatment with vehicle, AG-120, AZACITIDINE or the combination *in vivo*. **(C)** RRBS of I1DN cKit+ progenitor cells from mice treated for following 5 days of treatment with vehicle, AG-120, AZACITIDINE or the combination. Gene features associated with CpGs with sufficient coverage across all samples. **(D-F)** RNAseq of I1DN cKit+ progenitor cells from mice treated for following 5 days of treatment with vehicle, AG-120, AZACITIDINE or the combination. **(D)** MDS plot of top 500 most variably expressed genes. **(E)** Barcode plot demonstrating enrichment of AG-120 transcriptional signatures derived from previous experiments (see Figure 2) in mice engrafted with I1DN leukemias and treated with AG-120 once daily for 5 days. **(F)** Barcode plot demonstrating that AG-120 transcriptional signatures are enriched in the combination treatment. (E-F) ROAST analysis used to evaluate significance.

**SUPPLEMENTAL TABLES**

**Table S1: Graft-vs-host disease phenotype in AG-120 and dox-treated mice.**

**Table S2: Differentially expressed genes in I1DN progenitor cells sorted from mice treated with AG-120 or vehicle.**

**Table S3: Progenitor group specific differentially expressed genes induced by AG-120 treatment.**
